# Genetic variants of calcium and vitamin D metabolism in kidney stone disease

**DOI:** 10.1101/515882

**Authors:** Sarah A. Howles, Akira Wiberg, Michelle Goldsworthy, Asha L. Bayliss, Emily Grout, Chizu Tanikawa, Yoichiro Kamatani, Chikashi Terao, Atsushi Takahashi, Michiaki Kubo, Koichi Matsuda, Rajesh V. Thakker, Benjamin W. Turney, Dominic Furniss

## Abstract

Kidney stone disease (nephrolithiasis) is a major clinical and economic health burden^1,2^ with a heritability of ~45-60%^3^. To identify genetic variants associated with nephrolithiasis we performed genome-wide association studies (GWAS) and meta-analysis in British and Japanese populations, including 12,123 nephrolithiasis cases and 416,928 controls. Twenty loci associated with nephrolithiasis were identified, ten of which are novel. A novel *CYP24A1* locus is predicted to affect vitamin D metabolism and five loci, *DGKD, DGKH, WDR72, GPIC1*, and *BCR*, are predicted to influence calcium-sensing receptor (CaSR) signaling. In a validation cohort of nephrolithiasis patients the *CYP24A1*-associated locus correlated with serum calcium concentration and number of kidney stone episodes, and the *DGKD*-associated locus correlated with urinary calcium excretion. Moreover, DGKD knockdown impaired CaSR-signal transduction *in vitro*, an effect that was rectifiable with the calcimimetic cinacalcet. Our findings indicate that genotyping may inform risk of incident kidney stone disease prior to vitamin D supplementation and facilitate precision-medicine approaches, by targeting CaSR-signaling or vitamin D activation pathways in patients with recurrent kidney stones.

---

Kidney stones affect ~20% of men and ~10% of women by 70 years of age^1^ and commonly cause debilitating pain. The prevalence of this disorder is increasing and the United States is predicted to spend over $5 billion per year by 2030 on its treatment^2^. Unfortunately, up to 50% of individuals will experience a second kidney stone episode within 10 years of their initial presentation^4^ and recurrent stone disease is linked to renal function decline^5^.

Twin studies have reported a heritability of >45% and >50% for stone disease and hypercalciuria, respectively^3,6^ and a strong family history of urolithiasis, including a parent and a sibling, results in a standard incidence ratio (SIR) for stone formation of >50 in contrast to a SIR of 1.29 in spouses^7^. Three genome-wide association studies of nephrolithiasis have been published, identifying six loci associated with disease^8–10^. However, only two loci (*CLDN14* and *RGS14-SLC34A1*) have been replicated and no trans-ethnic studies have been undertaken. To increase understanding of the common genetic factors contributing to risk of nephrolithiasis, genome-wide association studies and meta-analysis were undertaken using UK Biobank and Biobank Japan resources^11,12^, integrating data from 12,123 stone formers and 416,928 controls.

Twenty genetic loci associating with nephrolithiasis were identified, 10 of which were initially identified from the UK Biobank discovery cohort (Supplementary Table 5) and another 10 from a subsequent trans-ethnic meta-analysis with Japanese GWAS summary statistics (Table 1, Fig. 1, and Supplementary Fig. 1)^13^. Fifty-four candidate genes were identified via *in silico* analysis of these 20 loci based on FUMA positional mapping, functional annotation, and biological plausibility^14^. MAGMA gene-property analysis implemented in FUMA revealed a striking overexpression of these genes in the kidney cortex; the GENE2FUNC tool demonstrated enrichment for gene ontologies associated with transmembrane ion transport, renal function, and calcium homeostasis, including “response to vitamin D” (Supplementary Fig. 2-3 and Table 7)^15^.

**Figure 1.**
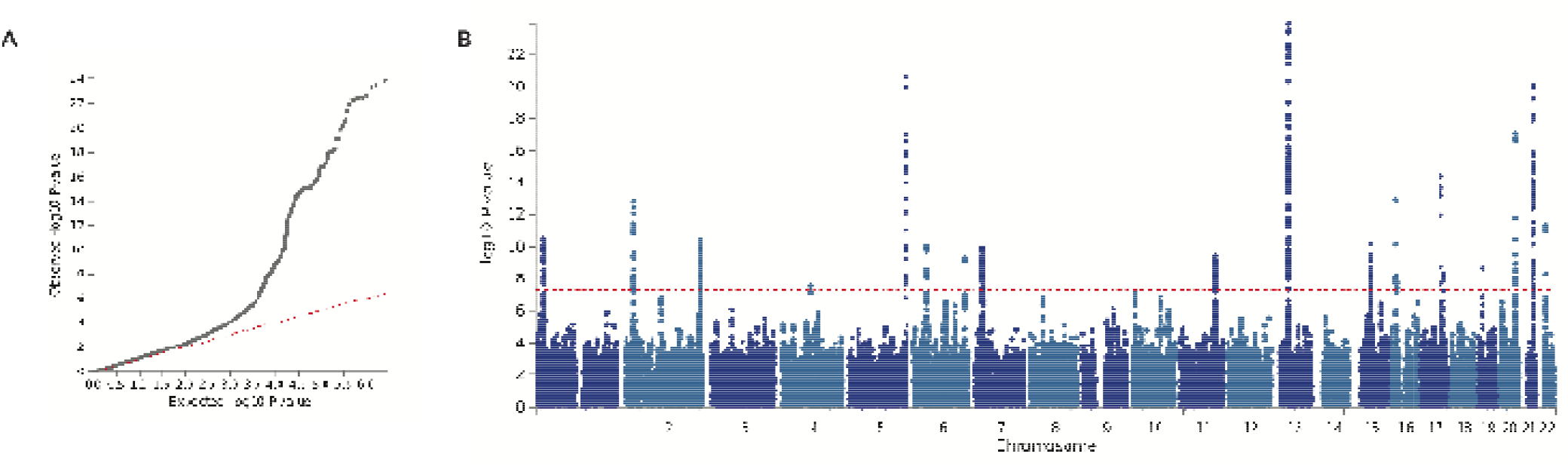
Results of trans-ethnic genome-wide association study in kidney stone disease. A trans-ethnic meta-analysis of kidney stone disease was performed for 12,123 patients with kidney stone disease and 416,928 controls from the UK Biobank and BioBank Japan. Panel A is a quantile-quantile plot of observed *vs*. expected p-values. The λ_GC_ demonstrated some inflation (1.0957), but the LD score regression (LDSC) intercept of 0.9997, with an attenuation ratio of 0.0075 indicated that the inflation was largely due to polygenicity and the large sample size. Panel B is a Manhattan plot showing the genome-wide p values (−log10) plotted against their respective positions on each of the autosomes. The horizontal red line shows the genome-wide significance threshold of 5.0x10^−8^.

**Table 1.**
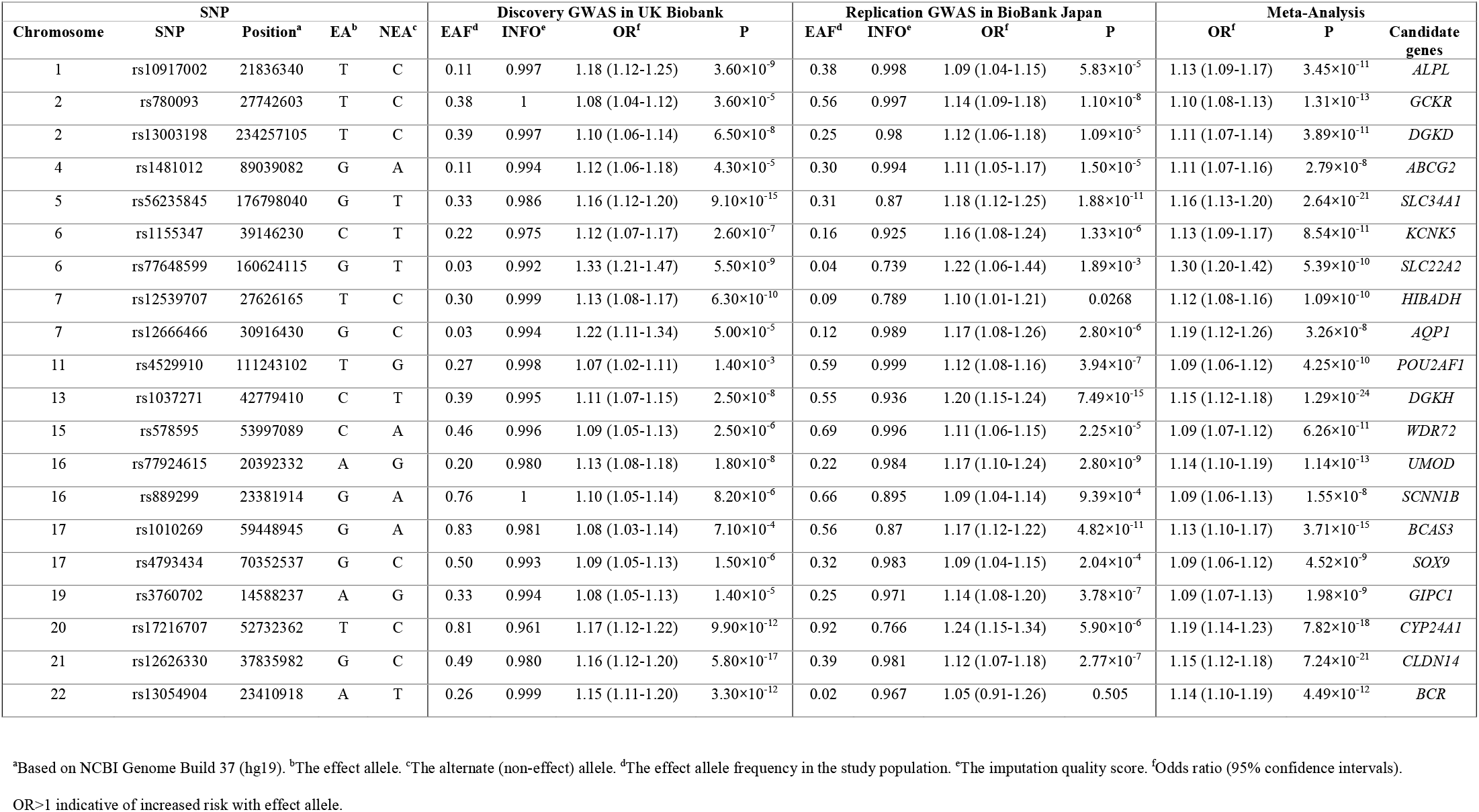
SNPs significantly associated with kidney stone disease at trans-ethnic meta-analysis.

Population Attributable Risk (PAR) was calculated at each locus using data from the UK Biobank population, and ranged from 1.3% to 22.3% (Supplementary Table 6), indicating the importance of these variants in the pathogenesis of nephrolithiasis at a population level. The highest PAR was identified with a *CYP24A1*-associated locus and five of the identified loci, with PAR scores between 5.9-7.9%, were predicted to influence CaSR-signaling. These loci were selected for further analysis.

rs17216707 is ~38kb upstream of *CYP24A1*, a gene encoding cytochrome P450 family 24 subfamily A member 1 (CYP24A1), an enzyme that metabolises active 1,25-dihydroxyvitamin D to inactive 24,25-dihydroxyvitamin D. Loss-of-function mutations in CYP24A1 cause autosomal recessive infantile hypercalcaemia type 1 (OMIM 126065)^16^. We postulated that the *CYP24A1* increased-risk allele associates with decreased CYP24A1 activity, leading to perturbations of calcium homeostasis and mimicking an attenuated form of infantile hypercalcaemia type 1. Therefore, associations of rs17216707 with serum calcium, phosphate, parathyroid hormone (PTH), and 25-hydroxyvitamin D concentrations, and urinary calcium excretion, and number of kidney stone episodes were sought in a validation cohort of kidney stone formers. 1,25-hydroxyvitamin D levels were unavailable. Reference ranges for 24-hour urinary calcium excretion differ for men and women^17^, thus associations with urinary calcium excretion were therefore examined separately.

Individuals homozygous for the *CYP24A1* increased-risk allele rs17216707 (T) had a significantly increased mean serum calcium concentration when compared to heterozygotes, consistent with a recessive effect (mean serum calcium 2.36mmol/l (TT) *vs*. 2.32mmol/l (TC)) (Table 2). rs17216707 (T) homozygotes had more kidney stone episodes than heterozygotes (mean number of stone episodes 4.0 (TT) *vs*. 2.4 (TC), p=0.0003) and there was a significant correlation across genotypes (TT *vs*. TC *vs*. CC) with number of stone episodes (p=0.0024)(Table 2). No correlation was found between rs17216707 genotype and serum phosphate, PTH, 25-hydroxyvitamin D concentration or urinary calcium excretion.

**Table 2:** Genotype-phenotype correlations in cohort of kidney stone formers.

| Variable | Normal range§ | CYP24A1 (rs17216707) |  |  | DGKD (rs838717) |  |  |
| --- | --- | --- | --- | --- | --- | --- | --- |
|  |  | TT | TC | CC | AA | AG | GG |
| **Serum** |  |  |  |  |  |  |  |
| Calcium (mmol/l) | 2.10-2.50 | **2.36±0.01\***<br>**(260)** | **2.32±0.01**<br>**(109)** | 2.34±0.02<br>(15) | 2.34±0.01<br>(107) | 2.35±0.01<br>(182) | 2.36±0.01<br>(95) |
| Phosphate (mmol) | 0.7-1.40 | 1.02±0.13<br>(274) | 1.02±0.02<br>(111) | 0.99±0.05<br>(14) | 1.03±0.02<br>(114) | 1.01±0.02<br>(193) | 1.02±0.02<br>(92) |
| Parathyroid hormone (pmol/l) | 1.3-7.6 | 5.06±0.18<br>(271) | 5.59±0.32<br>(107) | 5.27±0.64<br>(14) | 5.34±0.32<br>(108) | 5.38±0.23<br>(189) | 4.72±0.23<br>(95) |
| 25-hydroxy vitamin D (nmol/l) | >50 | 54.8±1.84<br>(227) | 50.6±2.60<br>(89) | 55.7±6.92<br>(10) | 56.1±3.13<br>(88) | 52.6±2.04<br>(157) | 53.2±2.90<br>(81) |
| **Urine** |  |  |  |  |  |  |  |
| Male patients<br>24hr calcium excretion (mmol) | <7.5 | 6.11±0.47<br>(77) | 4.82±0.55<br>(33) | 3.10±1.27<br>(3) | **4.54±0.45\***<br>**(33)** | 5.45±0.48<br>(57) | **7.27±0.91**<br>**(25)** |
| Female patients<br>24hr calcium excretion (mmol) | <6.2 | 4.85±0.53<br>(31) | 5.06±0.56<br>(15) | 4.27±1.69<br>(2) | 5.83±1.34<br>(9) | 4.92±0.46<br>(27) | 4.12±0.56<br>(12) |
| Number stone episodes | - | **4.0±0.4\***<br>**(287)** | **2.4±0.2**<br>**(19)** | 2.4±0.4<br>(14) | 3.5±0.41<br>(114) | 3.8±0.43<br>(208) | 2.8±0.26<br>(98) |

Numbers of stone forming patients included in analysis are shown in parentheses. Serum calcium values are albumin-adjusted. All values are expressed as mean ±SEM. Students T-tests were used for comparisons between groups of parametric data, Mann-Whitney-U tests were used for comparison of non-parametric data (number of stone episodes). Anova tests was used for comparisons of multiple sets of parametric data, no significance was reached. Kruskal-Wallis tests were used for comparisons of multiple sets of non-parametric data (number of stone episodes), significance at p=0.0024 was reached for *CYP24A1* locus correlations. \*Denotes significance on comparison to bold cohort within group at Bonferroni corrected threshold of $p < 0.05/7 = 0.007$ . §Normal ranges are from Nesbit et al<sup>26</sup> and Curhan et al<sup>17</sup>.

These findings support our hypothesis that the rs17216707 increased-risk allele is associated with a relative hypercalcaemia and reduced activity of the 24-hydroxylase enzyme and highlight the role of vitamin D catabolism in kidney stone formation. Patients with loss-of-function CYP24A1 mutations have been successfully treated with inhibitors of vitamin D synthesis including fluconazole, similar therapies may be useful in rs17216707 (TT) recurrent kidney stone formers^18^. Furthermore, vitamin D supplementation in patients with biallelic loss-of-function mutations in CYP24A1 cause nephrocalcinosis^16^. We predict that rs17216707 (TT) individuals may similarly display an increased sensitivity to vitamin D. The National Institute of Clinical Excellence (NICE) recommends that all adults living in the UK should take daily vitamin D supplementation. However, our findings suggest that supplementation may put individuals homozygous for the *CYP24A1* increased-risk allele at risk of kidney stones.

A previously reported association between the CaSR-associated intronic SNP rs7627468 and nephrolithiasis was confirmed (p=3.5x10^−5^)^9^. In addition, five of the identified loci are linked to genes that are predicted to influence CaSR signaling. rs13003198 is ~6kb upstream of *DGKD*, encoding diacylglycerol kinase delta (DGKD); rs1037271 is an intronic variant in *DGKH*, encoding diacylglycerol kinase eta (DGKH). DGKD and DGKH phosphorylate diacylglcerol, a component of the intracellular CaSR-signaling pathway inducing CaSR-mediated membrane ruffling and activating protein kinase C signaling cascades including MAPK^19,20^(Fig. 2). rs578595 is an intronic variant in *WDR72* encoding WD repeat domain 72 (WDR72) and rs3760702 is ~300bp upstream of *GIPC1* that encodes Regulator of G-protein signaling 19 Interacting Protein 1 (GIPC1). Both WDR72 and GIPC1 are thought to play a role in clathrin-mediated endocytosis, a process central to sustained intracellular CaSR signaling^20–23^(Fig. 2). rs13054904 is ~110kb upstream of *BCR*, encoding a RAC1 (Rac Family Small GTPase 1) GTPase-activating protein known as Breakpoint Cluster Region (BCR)^24^. RAC1 activation is induced by CaSR ligand binding and mediates CaSR-induced membrane ruffling^19^(Fig. 2).

**Figure 2.**
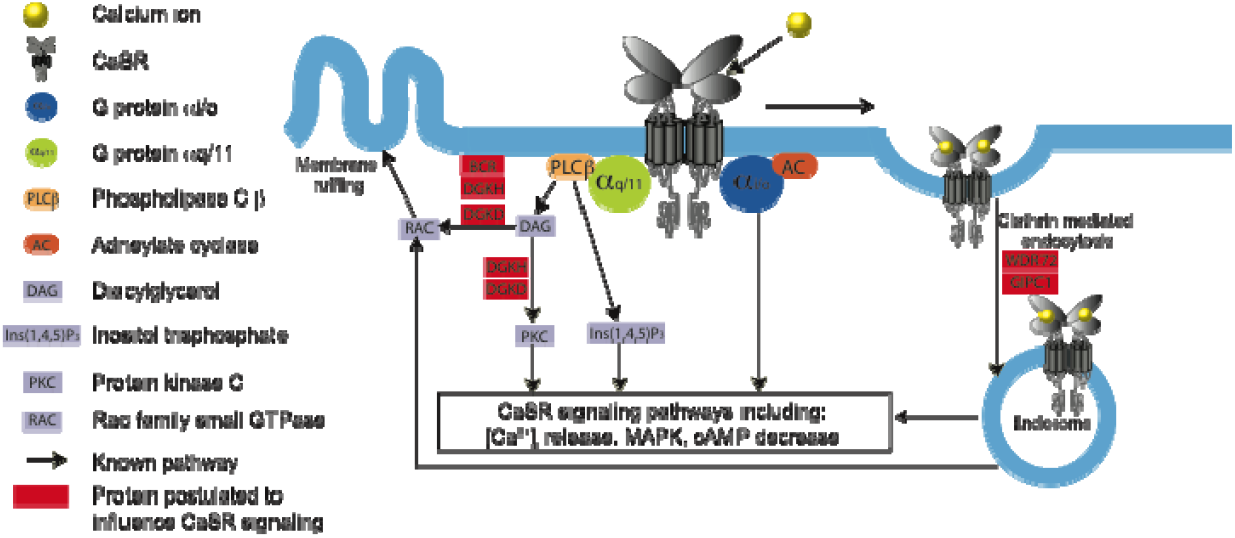
Schematic model for CaSR signaling. Ligand binding of calcium ions (yellow) by the G protein coupled receptor CaSR (gray) results in G protein-dependent stimulation via Gα_q/11_ (green) or Gα_i/o_ (blue) causing stimulation of intracellular signaling pathways including intracellular calcium ([Ca^2+^]_i_) release, MAPK stimulation or cAMP reduction. Gα_q/11_ signals via inositol 1,4,5-trisphosphate (IP3) and diacylglycerol (DAG). DAG leads to protein kinase C (PKC) stimulation along with RAC activation, which results in membrane ruffling. Following calcium ion binding the CaSR is internalized via clathrin-mediated endocytosis where signaling continues via the endosome. Proteins postulated to influence CaSR-signaling and their potential sites of action are shown in red.

**Figure 3.**
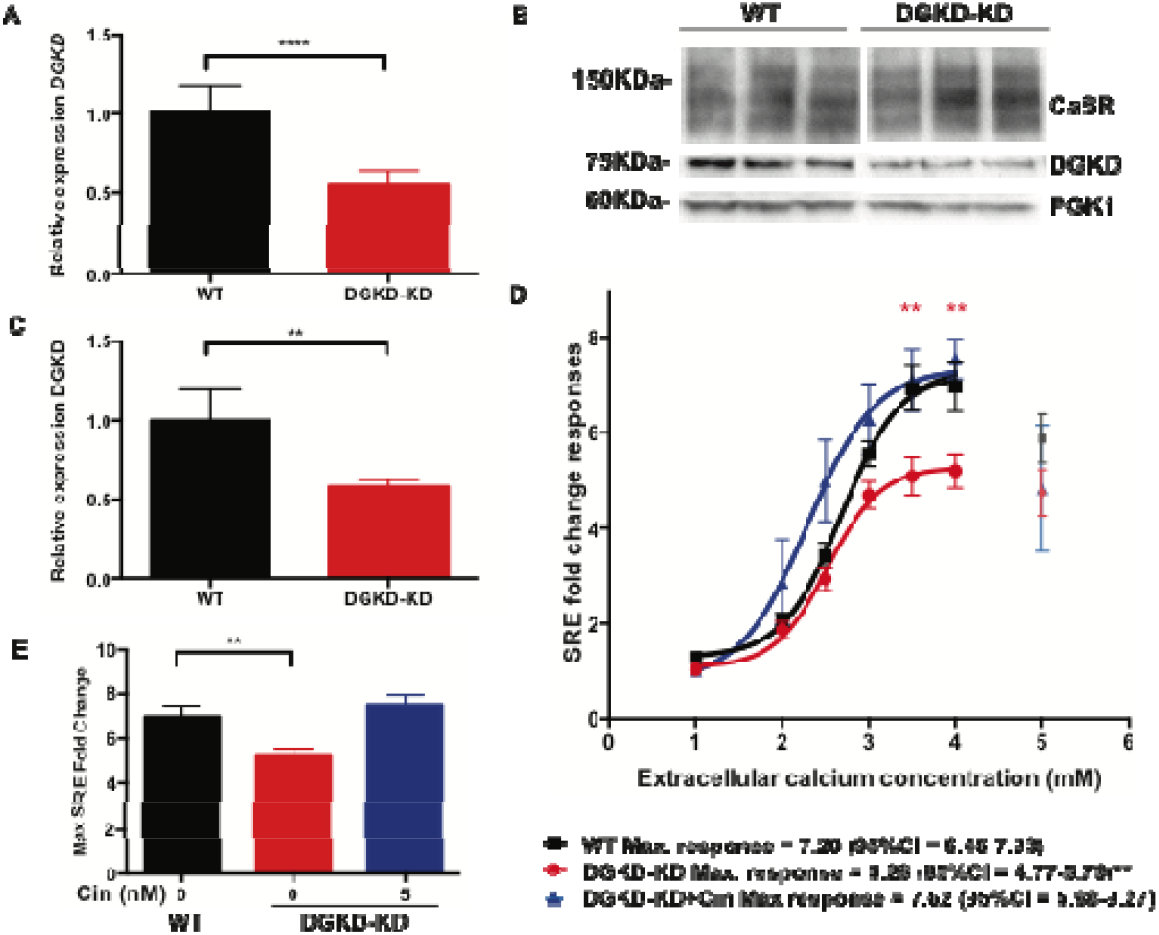
CaSR-mediated SRE responses following DGKD knockdown and effect of cinacalcet treatment in HEK-CaSR-SRE cells. Panel A shows relative expression of *DGKD*, as assessed by quantitative real-time PCR of HEK-CaSR-SRE cells treated with scrambled (WT) or *DGKD* (DGKD-KD) siRNA and used for SRE experiments. Samples were normalized to a geometric mean of four housekeeper genes: *PGK1, GAPDH, TUB1A, CDNK1B*. n=8. Panel B shows a representative western blot of lysates from HEK-CaSR cells treated with scrambled or *DGKD* siRNA and used for SRE experiments. PGK1 was used as a loading control. Panel C shows the relative expression of DGKD, as assessed by densitometry of western blots from cells treated with scrambled or *DGKD* siRNA demonstrating a ~50% reduction in expression of DGKD following treatment with *DGKD* siRNA. Samples were normalized to PGK1. All cells expressed CaSR. n=6. Panel D shows SRE responses of HEK-CaSR-SRE cells in response to changes in extracellular calcium concentration. Cells were treated with scrambled (WT) or *DGKD* (DGKD-KD) siRNA. The responses ± SEM are shown for 8 independent transfections for WT and DGKD-KD cells and 4 independent transfections for DGKD-KD + 5nM cinacalcet cells. Treatment with *DGKD* siRNA led to a reduction in maximal response (red line) compared to cells treated with scrambled siRNA (black line). This loss-of-function could be rectified by treatment with 5nM cinacalcet (blue line). Post desensitization points were not included in the analysis (grey, light red, and light blue). Panel E demonstrates the mean maximal responses with SEM of cells treated with scrambled siRNA (WT, black), *DGKD* siRNA (DGKD-KD, red) and *DGKD* siRNA incubated with 5nM cinacalcet (blue). Statistical comparisons of maximal response were undertaken using F test. Students T tests were used to compare relative expression. Two-way ANOVA was used to compare points on dose response curve with reference to WT. Data are shown as mean ±SEM with **p<0.01, ****p<0.0001.

The CaSR plays a central role in calcium homeostasis, increasing renal calcium reabsorption and stimulating PTH release to enhance bone resorption, urinary calcium reabsorption, and renal synthesis of 1,25-dihydroxyvitamin D^25^. Gain-of-function mutations in components of the CaSR-signaling pathway result in autosomal dominant hypocalcaemia (ADH, OMIM 601198, 615361), which is associated with hypercalciuria in ~10% of individuals^26,27^. ADH-associated mutations result in a gain-of-function in CaSR-intracellular signaling *in vitro* via pathways including intra-cellular calcium ions or MAPK^26,28,29^. We hypothesized that the nephrolithiasis-associated loci linked to CaSR-signaling associate with enhanced CaSR signal transduction resulting in a biochemical phenotype mimicking an attenuated form of ADH. *DGKD* and *DGKH* were selected for further analysis based on their PAR scores and the potential to influence both membrane ruffling and MAPK CaSR-signaling pathways (Fig.2).

The *DGKD* increased-risk allele rs838717 (G) (top *DGKD*-associated SNP in the UK Biobank GWAS, Supplementary Table 4, linkage disequilibrium with rs13003198 r^2^=0.53) associated with increased 24-hour urinary calcium excretion in male stone formers (mean 24-hour urinary calcium excretion 7.27mmol (GG) *vs*. 4.54mmol (AA), p=0.0055) (Table 2) consistent with enhanced CaSR-signal transduction. No association of the *DGKD* increased-risk allele rs838717 (G) with urinary calcium excretion was identified in female stone formers, which is likely due to a lack of power as a result of the small sample size, however it is interesting to note that heritability of stone disease is lower in women than in men^3^. Furthermore, no correlations were observed between genotype and serum calcium, phosphate, PTH, 25-hydroxyvitamin D concentrations or number of stone episodes despite hypocalcemia being the prominent phenotype in ADH patients. This may relate to differential tissue expression patterns of DGKD or the specificity of DGKD effects on intracellular CaSR-signaling pathways.

The *DGKH* increased-risk allele rs1170174 (A) (top *DGKH*-associated SNP in the UK Biobank GWAS, Supplementary Table 4, linkage disequilibrium with rs13003198 r^2^=0.3) did not associate with biochemical phenotype or stone recurrence. However the sample size of homozygotes was small (AA, n=6) and prior to Bonnferonni correction, a suggestive association was detected was detected with urinary calcium excretion in male stone formers (mean 24-hour urinary calcium excretion 8.14mmol (AA) *vs*. 5.09mmol (GG), p=0.0503).

To investigate the role of DGKD in CaSR-signalling, HEK-CaSR-SRE cells were treated with scrambled or *DGKD* targeted siRNA and intracellular MAPK responses to alterations in extracellular calcium concentration assessed. SRE responses were significantly decreased in cells with reduced DGKD expression (DGKD-KD) when compared to cells with baseline DGKD expression (WT) (maximal response DGKD-KD=5.28 fold change, 95% confidence interval (CI)=4.77-5.79 *vs*. WT=7.20 fold change, 95% CI=6.46-7.93, p=0.0065). Cinacalcet, rectified this loss-of-function (DGKD-KD+5nM cinacalcet, maximal response=7.62 fold change, 95% CI=5.98-9.27)(Fig. 2A-E). These findings provide evidence that DGKD influences CaSR-mediated signal transduction and suggest that the *DGKD* increased-risk allele may associate with a relative increase in DGKD expression thereby enhancing CaSR-mediated signal transduction. Calcilytics, including NPS-2143 and ronacaleret, rectify enhanced CaSR-mediated signaling *in vitro* and biochemical phenotypes in mouse models of ADH^29–31^ and therefore may represent novel, targeted therapies for recurrent stone formers carrying CaSR-associated increased-risk alleles.

In conclusion, this study has identified 20 loci linked to kidney stone formation, 10 of which are novel, and revealed the importance of vitamin D metabolism pathways and enhanced CaSR-signaling in the pathogenesis of nephrolithiasis. Our findings suggest a role for genetic testing to identify individuals in whom vitamin D supplementation should be used with caution and to facilitate a precision-medicine approach for the treatment of recurrent kidney stone disease, whereby targeting of the CaSR-signaling or vitamin D metabolism pathways may be beneficial in the treatment of a subset of patients with nephrolithiasis.

## Acknowledgements

This work was supported by grants from Kidney Research UK (RP_030_20180306) to S.A.H., A.W., M.G., B.W.T., and D.F, National Institute for Health Research (N.I.H.R) Oxford Biomedical Research Centre to R.V.T, and the Wellcome Trust (204826/z/16/z) to S.A.H, and M.G. S.A.H. is a N.I.H.R Academic Clinical Lecturer. A.W is a MRC Clinical Research Training Fellow. R.V.T. has Senior Investigator Awards from the Wellcome Trust (106995/z/15/z) and N.I.H.R. (NF-SI-0514-10091). We acknowledge the contribution to this study made by the Oxford Centre for Histopathology Research and the Oxford Radcliffe Biobank, which are supported by the NIHR Oxford Biomedical Research Centre.

## Author Contributions

S.A.H., A.W., B.W.T., and D.F. designed this study. S.A.H, A.W., M.G., A.L.B., E.G., C.Ta., Y.K., C.Te., A.T., M.K., K.M., and B.W.T acquired data. A.W. and D.F. carried out genetic association analysis. S.A.H, M.G., and A.L.B undertook *in vitro* studies. S.A.H., A.W., M.G, C.Ta., Y.K., C.Te., A.T., M.K., K.M., R.V.T, B.W.T., and D.F analysed and interpreted data. S.A.H., A.W., R.V.T., B.W.T., and D.F wrote the first draft of the manuscript. All other co-authors participated in the preparation of the manuscript by reading and commenting on the draft prior to submission.

## Completing Financial Interests

The authors declare no competing financial interests.

## Online Methods

### Study participants

#### UK Biobank

For the UK-based GWAS, the UK Biobank resource was utilised, a prospective cohort study of ~500,000 individuals from the UK, aged between 40-69, who have had whole-genome genotyping undertaken, and have allowed linkage of these data with their medical records^1,2^. ICD-10 and OPCS codes were used to identify individuals with a history of nephrolithiasis (Supplementary Table 1). Patients were excluded if they were recorded to have a disorder of calcium homeostasis, malabsorption, or other condition known to predispose to kidney stone disease (Supplementary Table 2). Following quality control (QC) 6,536 UK Biobank participants were identified as cases and 388,508 individuals as controls (Supplementary Table 3).

#### Japanese patients

Diagnosis of nephrolithiasis was confirmed by enrolling physicians; patients with bladder stones were excluded. DNA samples of 5,587 nephrolithiasis patients were obtained from BioBank Japan^3^. Controls (28,870 individuals) were identified from four population-based cohorts, including the JPHC (Japan Public Health Center)-based prospective study^4^, the J-MICC (Japan Multi-Institutional Collaborative Cohort) study^5^, IMM (Iwate Tohoku Medical Megabank Organization) and ToMMo (Tohoku Medical Megabank Organization)^6^.

#### Validation cohort

Patients attending the Oxford University Hospitals NHS Foundation Trust for treatment of kidney stones were enrolled into the Oxford University Hospitals NHS Foundation Trust Biobank of Kidney Stone Formers, following informed consent. Clinical data including urological history, past medical history and family history was recorded along with details of serum and urinary biochemistry. Whole blood and urine samples were stored at −80°C.

### Ethical approval

UK Biobank has approval from the North West Multi-Centre Research Ethics Committee (11/NW/0382), and this study (“Epidemiology of Kidney Stone Disease”) has UK Biobank study ID 885. Ethical committees at each Japanese institute approved the project. Collection of clinical data and biological samples from kidney stone patients attending the Oxford University Hospitals NHS Foundation Trust was approved under the Oxford Radcliffe Biobank research tissue bank ethics (09/H0606/5+5). All patients provided written informed consent.

### Genotyping

#### UK Biobank

The genotyping, QC and imputation methodology employed by UK Biobank are described in detail elsewhere^7^. Briefly, UK Biobank contains genotypes of 488,377 participants who were genotyped on two very similar genotyping arrays: UK BiLEVE Axiom Array (807,411 markers; 49,950 participants), and UK Biobank Axiom Array (825,927 markers; 438,427 participants). The two arrays are very similar, sharing approximately 95% of marker content. Genotypes were called from the array intensity data, in 106 batches of approximately 4700 samples each using a custom genotype-calling pipeline.

#### Japanese samples

Samples were genotyped with Illumina HumanOmniExpressExome BeadChip or a combination of the Illumina HumanOmniExpress and HumanExome BeadChips^8,9^.

#### Validation cohort

DNA was extracted from whole blood samples using the Maxwell 16 Tissue DNA purification kit (Promega). SNP genotyping was undertaken using TaqMan SNP Genotyping Assays (ThermoFisher) and Type-it^®^ Fast SNP Probe PCR Kit (Qiagen).

### Quality Control

#### UK Biobank

Quality Control (QC) was performed using PLINK^10^ v1.9 and R v3.3.1. All SNPs with a call rate <90% were removed, accounting for the two different genotyping platforms used to genotype the individuals. Sample-level QC was undertaken and individuals excluded with one or more of the following: (1) call rate <98%, (2) discrepancy between genetically inferred sex (Data Field 22001) and self-reported sex (Data Field 31), or individuals with sex chromosome aneuploidy (Data Field 22019), (3) heterozygosity >3 S.D. from the mean (calculated using UK Biobank’s PCA-adjusted heterozygosity values, Data Field 20004). Individuals were then excluded who were not flagged by UK Biobank as having white British ancestry (on the basis of principal component analysis and self-reporting as “British” – Data Field 22006). Data was merged with publicly available data from the 1000 Genomes Project^5,11^ and principal components analysis (PCA) performed using flashpca^12^ to confirm that the white British ancestry individuals from UK Biobank overlapped with the “GBR” individuals from the 1000 Genomes Project. BOLT-LMM was used in analysis and therefore there were no sample exclusions based on relatedness^13^. in total, 86,693 individuals were excluded based on the above criteria. SNP-level QC was performed by excluding SNPs with Hardy-Weinberg equilibrium (HWE) p<10^−4^, <98% call rate, and minor allele frequency (MAF)<1%. Overall, 237,245 SNPs were excluded in total. Finally, six individuals were excluded who harboured an abnormal number of SNPs with a minor allele count of 1, or were visual outliers when autosomal heterozygosity was plotted against call rate. This resulted in a final dataset of 401,667 individuals and 547,011 SNPs. Following QC, individuals were excluded who were recorded to have a disorder of calcium homeostasis, malabsorption, or other condition known to predispose to kidney stone disease (Supplementary Table 2). Subsequent case ascertainment was performed using the list of ICD-10 and OPCS codes for kidney and ureteric stones (Supplementary Table 1).

#### Japanese Samples

Samples were excluded if (i) call rate□ was < 0.98, (ii) they were from closely related individuals identified by identity-by-descent analysis, (iii) the samples were sex-mismatched with a lack of information, or (iv) they were non–East Asian outliers identified by principal component analysis of the studied samples and the three major reference populations (Africans, Europeans, and East Asians) in the International HapMap Project^14^. Standard quality-control criteria for variants were applied, excluding those with (i) SNP call rate < 0.99, (ii) minor allele frequency < 1%, and (iii) Hardy–Weinberg equilibrium P value < 1.0 × 10^−6^.

### Imputation

#### UK Biobank data

UK Biobank’s method of phasing and imputation of SNPs is described in detail elsewhere^7^. Briefly, phasing on the autosomes was performed using SHAPEIT3^15^, using the 1000 Genomes Phase 3 dataset as a reference panel. For imputation, both the HRC (Haplotype Reference Consortium) reference panel^16^ and a merged UK10K / 1000 Genomes Phase 3 panel were used. This resulted in a dataset with 92,693,895 autosomal SNPs, short indels and large structural variants. Imputation files were released in the BGEN (v1.2) file format.

#### Japanese data

Genotypes were prephased with MACH^17^ and dosages imputed with minimac and the 1000 Genomes Project Phase 1 (version 3) East Asian reference haplotypes^11^.

### Association Analysis

#### UK Biobank data

Genome-wide association analysis was undertaken across 547,011 genotyped SNPs and ~8.4 million imputed SNPs with MAF≥0.01 and Info Score≥0.9, using a linear mixed non-infinitesimal model implemented in BOLT-LMM v2.3^18^. A reference genetic map file for hg19 and a reference linkage disequilibrium (LD) score file for European-ancestry individuals included in the BOLT-LMM package in the analysis was used. Three covariates were used in the association study: genetic sex, age, and the genotyping platform (to account for array effects). The LD score regression intercept^11^ of 0.9997 with an attenuation ratio of 0.0075 indicated minimal inflation when adjusted for the large sample size. Conditional analysis at each associated locus was performed by conditioning on the allelic dosage (calculated using QCTOOL v2) of the most significantly associated SNP at each locus.

#### Japanese data

GWAS was conducted using a logistic regression model by incorporating age, sex, and the top 10 principal components as covariates.

#### Trans-ethnic Meta-analysis

Trans-ethnic meta-analysis was performed using the summary statistics from the UK and Japanese GWAS data sets (12,123 cases and 416,928 controls). An imputation quality score (RSQR) threshold of >0.5 was applied to the Japanese GWAS SNPs^19^ prior to performing a fixed-effects meta-analysis using GWAMA^20^, using ~5 million SNPs common to both GWAS datasets. Quantile-quantile and Manhattan plots were created using FUMA^21^.

### *In Silico* Analyses

The summary statistics from the GWAS meta-analysis were analysed in FUMA^21^ v1.3.3c, selecting UK Biobank Release 2 (White British) as the population reference panel (as the vast majority of individuals in the meta-analysis are of British rather than Japanese ethnicity). Functionally annotated SNPs were mapped to genes based on genomic position and annotations obtained from ANNOVAR, using positional mapping in FUMA (Supplementary Table 7)^22^. MAGMA (implemented in FUMA) was used to perform a gene-property analysis in order to identify particular tissue types relevant to kidney stones. This analysis determines if tissue-specific differential expression levels are predictive of the association of a gene with kidney stones, across 53 different tissues taken from the GTEx v7 database^23^ (Supplementary Figure 2).

In order to gain insight into the biological pathways implicated by the FUMA-prioritised genes, a gene set analysis was implemented using the GENE2FUNC tool in FUMA using the 54 positionally mapped genes with unique Entrez IDs and gene symbols. The following parameters were applied: Benjamini-Hochberg false discovery rate (FDR) for multiple testing correction, adjusted p-value cut-off = 0.0025, minimum number of overlapped genes = 2, GTEx v7 RNA-Seq expression data. Hypergeometric tests were performed to test if genes of interest are overrepresented in any of the pre-defined gene sets in GO biological processes (MsigDB v6.1) (Supplementary Figure 3).

### Population Attributable Risk

To assess the impact of an identified allele of interest from the meta-analysis with risk of kidney stone disease within the UK Biobank population, Population Attributable Risk (PAR) was calculated for SNPs of interest using the following equations^24^ with individual-level genotype data from the UK Biobank:

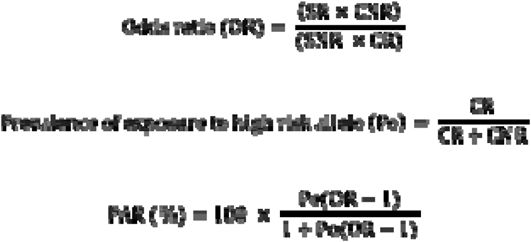

Where:

SR = Stone formers with at least one increased-risk allele
SNR = Stone formers with no risk allele
CR = Control patients with at least one increased-risk allele
CNR = Control patients with no risk allele

Dosages of risk alleles for all individuals were computed using QCTOOL v2. As the majority of SNPs were imputed rather than genotyped, allelic dosages were not always discrete (i.e. 0, 1 or 2). In such circumstances, an allelic dosage threshold of ≥ 0.9 was applied in order to consider an individual to possess at least one copy of the risk allele.

### Genotype-Phenotype Correlations

Associations were sought between genotype and serum calcium (albumin adjusted), phosphate, parathyroid hormone, and 25-hydroxyvitamin D, and urinary calcium excretion and number of stone episodes. Patients were excluded from inclusion in genotype-phenotype correlations if they were known to have a disorder of calcium homeostasis, malabsorption, or other condition known to predispose to kidney stone disease.

### Functional characterisation of the Calcium-Sensing Receptor Pathway

#### Cell culture and transfection

Functional studies were undertaken using HEK293 cells that had been transfected to stably express calcium-sensing receptors (CaSRs) and luciferase under the control of a serum-response element (SRE) (HEK-CaSR-SRE cells). Cells were transfected with scrambled or *DGKD* siRNA (Qiagen) using lipofectamine RNAiMAX (Thermo Fisher Scientific) 72 hours before experiments. HEK-CaSR-SRE cells were maintained in DMEM-Glutamax media (Thermo Fisher Scientific) with 10% FBS (Gibco) and 400μg/ml geneticin (Thermo Fisher Scientific) and 200μg/ml hygromycin (Invitrogen) at 37°C, 5% CO_2_.

#### Confirmation of DGKD knockdown

Successful knockdown of DGKD was confirmed via quantitative reverse transcriptase PCR (qRT-PCR) and western blot analyses. qRT-PCR analyses were performed in quadruplicate using Power SYBR Green Cells-to-CT™ Kit (Life Technologies), *DGKH, PGK1, GAPDH, TUB1A, CDNK1B* specific primers (Qiagen), and a Rotor-Gene Q real-time cycler (Qiagen Inc, Valencia, CA). Samples were normalized to a geometric mean of four housekeeper genes: *PGK1, GAPDH, TUB1A, CDNK1B*. Western blot analyses were undertaken using anti-DGKD (SAB1300472; Sigma), anti-CaSR (5C10, ADD; ab19347; Abcam), and anti-PGK1 (ab154613; Abcam). The western blots were visualized using an Immuno-Star Western C kit (Bio-Rad) on a Bio-Rad Chemidoc XRS+ system and relative expression of DGKD was quantified by denistometry using ImageJ software.

#### Compounds

Cinacalcet (AMG-073 HCL) was obtained from Cambridge Bioscience (catalog CAY16042) and dissolved in DMSO prior to use in *in vitro* studies.

#### SRE Response Assays

At 60 hours post transfection, cells were incubated in 0.05% fetal bovine serum media with 0.45mM calcium for 12 hours. At 72 hours the media was changed to varying concentrations of extracellular calcium (0.1-5mM), with either 5nM cinacalcet or equivalent volume of DMSO, and the cells were incubated for a further 4 hours at 37°C. Cells were lysed and luciferase activity measured using Luciferase Assay System (Promega) on the PHERAstar microplate reader (BMG Labtech). Assays were performed in >4 biological replicates (independently transfected wells, performed on at least 4 different days). Nonlinear regression of concentration-response curves was performed with GraphPad Prism for determinations of maximal response and EC_50_ values.

### Statistics

For genotype-phenotype correlations, 2-tailed Student T Tests were used for parametric data. Mann-Whitney-U tests were used for comparison of non-parametric data (number of stone episodes). Anova tests were used for comparisons of multiple sets of parametric data. Kruskall-Wallis tests were used for comparisons of multiple sets of non-parametric data (number of stone episodes). Significance was defined as p<0.05 after Bonferroni correction for 7 tests on each set of data, thus p<0.05/7=0.007.

For *in vitro* studies statistical comparisons were made with 2-tailed Student T Tests and maximal responses were compared using the *F*-test^25^.

