## Supplementary appendix for "Genetic variants of calcium and vitamin D metabolism in kidney stone disease"

This appendix has been provided by the authors to give readers additional information about their work

**Table of Contents**

1	Tables	3

Table 1: Inclusion criteria for identification of stone forming individuals in UK Biobank population	3

Table 2: Exclusions for UK Biobank population	4

Table 3: UK Biobank Study Population	5

Table 4: Japanese Study population	6

Table 5. SNPs significantly associated with kidney stone disease in UK Biobank population	7

Table 6: Population Attributable Risk in the UK Biobank population	8

2	Figures	10

Figure 1	10

Figure 2	21

Figure 3	22

### 1 Tables

#### Table 1: Inclusion criteria for identification of stone forming individuals in UK Biobank population

| **ICD-10 Codes** | | **OPCS codes** | |
| --- | --- | --- | --- |
| N20.0 | Calculus of kidney | M061 | Open removal of calculus from kidney |
| N20.1 | Calculus of ureter | M091 | Endoscopic ultrasound fragmentation of calculus of kidney |
| N20.2 | Calculus of kidney with calculus of ureter | M092 | Endoscopic electrohydraulic shock wave fragmentation of calculus of kidney |
| N20.9 | Urinary calculus, unspecified | M093 | Endoscopic laser fragmentation of calculus of kidney |
| N23 | Unspecified Renal Colic | M094 | Endoscopic extraction of calculus of kidney |
|  |  | M098 | Other specified therapeutic endoscopic operations on calculus of kidney, |
|  |  | M099 | Unspecified therapeutic endoscopic operations on calculus of kidney |
|  |  | M141 | Extracorporeal shock wave lithotripsy of calculus of kidney |
|  |  | M148 | Other specified extracorporeal fragmentation of calculus of kidney |
|  |  | M149 | Unspecified extracorporeal fragmentation of calculus of kidney |
|  |  | M164 | Percutaneous nephrolithotomy |
|  |  | M271 | Ureteroscopic laser fragmentation of calculus of ureter |
|  |  | M272 | Ureteroscopic fragmentation of calculus of ureter |
|  |  | M273 | Ureteroscopic extraction of calculus of ureter |
|  |  | M281 | Endoscopic laser fragmentation of calculus of ureter |
|  |  | M282 | Endoscopic fragmentation of calculus of ureter |
|  |  | M283 | Endoscopic extraction of calculus of ureter |
|  |  | M284 | Endoscopic catheter drainage of calculus of ureter |
|  |  | M288 | Other specified other endoscopic removal of calculus from ureter |
|  |  | M289 | Unspecified other endoscopic removal of calculus from ureter |
|  |  | M311 | Extracorporeal shock wave lithotripsy of calculus of ureter |
|  |  | M318 | Other specified extracorporeal fragmentation of calculus of ureter |
|  |  | M319 | Unspecified extracorporeal fragmentation of calculus of ureter |
|  |  | M26.1 | Nephroscopic laser fragmentation of calculus of ureter |
|  |  | M26.2 | Nephroscopic fragmentation of calculus of ureter NEC |
|  |  | M26.3 | Nephroscopic extraction of calculus of ureter |
|  |  | M28.5 | Endoscopic drainage of calculus of ureter by dilation of ureter |
|  |  | M28.8 | Other specified other endoscopic removal of calculus from ureter |

#### Table 2: Exclusions for UK Biobank population

| **ICD-10 Codes** | | **OPCS codes** | |
| --- | --- | --- | --- |
| E26.8 | Bartter syndrome | M39.1 | Open removal of calculus from bladder |
| E72.0 | Disorders of amino acid transport | M44.2 | Endoscopic extraction of calculus of bladder |
| E21.0 | Hyperparathyroidism | M67.4 | Endoscopic removal of calculus from prostate |
| E21.1 | Hyperparathyroidism | M75.8 | Open extraction of calculus from urethra |
| E21.2 | Hyperparathyroidism | G27.1 | Gastric bypass surgery |
| E21.3 | Hyperparathyroidism | G27.2 | Gastric bypass surgery |
| Q61.5 | Medullary sponge kidney | G27.3 | Gastric bypass surgery |
| N25.8 | Type 1 renal tubular acidosis | G27.4 | Gastric bypass surgery |
| K50 | Inflammatory bowel disease | G27.5 | Gastric bypass surgery |
| K51 | Inflammatory bowel disease | G27.8 | Gastric bypass surgery |
| K91.2 | Postsurgical malabsorption | G27.2 | Gastric bypass surgery |
| Q62 | Congential obstructive defects of the renal pelvis and malfomations of the ureter | G28.1 | Gastric bypass surgery |
| E83.31 | Hereditrary hypophosphatemic rickets with hypercalciruria and nephrolithiasis, osteoporosis and hypophosphatemia | G28.2  G28.3 | Gastric bypass surgery  Gastric bypass surgery |
| E83.42 | Familial hypomagnesemia with hypercalciuria and nephrocalcinosis and Familial hypomagnesemia with hypercalciuria and nephrocalcinosis with ocular abnormalities | G28.4  G28.5 | Gastric bypass surgery  Gastric bypass surgery |
| E74.8 | Oxaluria and oxalosis | G28.8  G28.9 | Gastric bypass surgery  Gastric bypass surgery |
| N21.0 | Calculus in bladder | G31.1 | Gastric bypass surgery |
| N21.1 | Calculus in urethra | G31.2 | Gastric bypass surgery |
| N21.8 | Other lower urinary tract calculus | G31.3 | Gastric bypass surgery |
| N21.9 | Calculus of the lower urinary tract | G31.4 | Gastric bypass surgery |
|  |  | G31.8 | Gastric bypass surgery |
|  |  | G31.9 | Gastric bypass surgery |
|  |  | G31.0 | Gastric bypass surgery |
|  |  | G32.1 | Gastric bypass surgery |
|  |  | G32.2 | Gastric bypass surgery |
|  |  | G32.3 | Gastric bypass surgery |
|  |  | G32.4 | Gastric bypass surgery |
|  |  | G32.8 | Gastric bypass surgery |
|  |  | G32.9 | Gastric bypass surgery |
|  |  | G32.0 | Gastric bypass surgery |
|  |  | G33.1 | Gastric bypass surgery |
|  |  | G33.2 | Gastric bypass surgery |
|  |  | G33.3 | Gastric bypass surgery |
|  |  | G33.6 | Gastric bypass surgery |
|  |  | G33.8 | Gastric bypass surgery |
|  |  | G33.9 | Gastric bypass surgery |
|  |  | G33.0 | Gastric bypass surgery |

#### Table 3: UK Biobank Study Population

| **Cohort** | **Number of Samples** | **Female (%)** | **Male (%)** | **Mean Age (SD)** |
| --- | --- | --- | --- | --- |
| Kidney stone | 6,536 | 33.3 | 66.7 | 67.9 (7.63) |
| Control | 388,508 | 54.5 | 45.5 | 66.8 (8.01) |

#### Table 4: Japanese Study population

| **Cohort** | **Source** | **Genotyping Platform** | **Number of Samples** | **Female (%)** | **Male (%)** | **Mean Age (SD)** |
| --- | --- | --- | --- | --- | --- | --- |
| Kidney stone | BBJ | HumanOmniExpressExome or HumanOmniExpress and HumanExome | 5,587 | 1,333 (23.9) | 4,254 (76.1) | 53.7 (13.8) |
| Control | JPHC  J-MICC  ToMMo | HumanOmniExpressExome | 28,870 | 17,492 (60.6) | 11,378 (43.7) | 56.3 (10.0) |

BBJ: Biobank Japan, JPHC: Japan Public Health Centre-based Prospective Study; J-MICC: Japan Multi-Institutional Collaborative Cohort Study; ToMMo: Tohoku Medial Megabank Organisation

#### Table 5. SNPs significantly associated with kidney stone disease in UK Biobank population

| **Chromosome** | **Position^a^** | **rsID** | **Effect Allele** | **Non-Effect Allele** | **EAF^b^** | **INFO score** | **OR (95% CI)** | **P** | **Candidate Gene** |
| --- | --- | --- | --- | --- | --- | --- | --- | --- | --- |
| 1 | 21836934 | rs6703976 | T | C | 0.11 | 0.998 | 1.18 (1.12-1.25) | 3.7**×**10^-9^ | *ALPL* |
| 1 | 21893344 | rs1256332 | A | C | 0.16 | 0.995 | 1.17 (1.12-1.23) | 6.2**×**10^-11^ | *ALPL* |
| 2 | 234296444 | rs838717 | G | A | 0.43 | 0.995 | 1.11 (1.07-1.15) | 1.6**×**10^-8^ | *DGKD* |
| 5 | 17679999 | rs10051765 | C | T | 0.33 | 0.995 | 1.16 (1.11-1.20) | 7.8**×**10^-15^ | *SLC34A1* |
| 6 | 160619918 | rs28495851 | C | A | 0.03 | 0.989 | 1.35 (1.22-1.50) | 4.7**×**10^-9^ | *SLC22A2* |
| 7 | 27653207 | rs7790498 | A | G | 0.30 | 0.999 | 1.13 (1.08-1.17) | 2.9**×**10^-10^ | *HIBADH* |
| 13 | 42688211 | rs1170174 | A | G | 0.18 | 0.988 | 1.16 (1.11-1.22) | 4.5**×**10^-11^ | *DGKH* |
| 16 | 20392332 | rs77924615 | A | G | 0.20 | 0.98 | 1.13 (1.09-1.18) | 1.8**×**10^-8^ | *UMOD* |
| 20 | 52732362 | rs17216707 | T | C | 0.81 | 0.961 | 1.17 (1.12-1.22) | 9.9**×**10^-12^ | *CYP24A1* |
| 21 | 37818871 | rs2776288 | A | G | 0.63 | 0.988 | 1.18 (1.14-1.22) | 5.7**×**10^-19^ | *CLDN14* |
| 22 | 23410918 | rs13054904 | A | T | 0.26 | 0.999 | 1.15 (1.11-1.20) | 3.3**×**10^-12^ | *BCR* |

^a^Based on NCBI Genome Build 37 (hg19). ^b^The effect allele frequency in kidney stone formers. Two independent signals were identified at the *ALPL* locus.

#### Table 6: Population Attributable Risk in the UK Biobank population

| Chromosome | rsID | Candidate Genes | PAR (%) |
| --- | --- | --- | --- |
| **1** | rs10917002 | *ALPL* | 3.27 |
| **2** | rs780093 | *GCKR* | 5.24 |
| **2** | rs13003198 | *DGKD* | 7.02 |
| **4** | rs1481012 | *ABCG2* | 2.53 |
| **5** | rs56235845 | *SLC34A1* | 9.08 |
| **6** | rs1155347 | *KCNK5* | 4.84 |
| **6** | rs77648599 | *SLC22A2* | 1.91 |
| **7** | rs12539707 | *HIBADH* | 7.52 |
| **7** | rs12666466 | *AQP1* | 1.30 |
| **11** | rs4529910 | *POU2AF1* | 3.86 |
| **13** | rs1037271 | *DGKH* | 7.92 |
| **15** | rs578595 | *WDR72* | 6.32 |
| **16** | rs77924615 | *UMOD* | 4.91 |
| **16** | rs889299 | *SCNN1B* | 13.52 |
| **17** | rs1010269 | *BCAS3* | 14.10 |
| **17** | rs4793434 | *SOX9* | 6.79 |
| **19** | rs3760702 | *GIPC1* | 5.92 |
| **20** | rs17216707 | *CYP24A1* | 22.31 |
| **21** | rs12626330 | *CLDN14* | 13.57 |
| **22** | rs13054904 | *BCR* | 6.62 |

**Table 7. Genes implicated in FUMA positional mapping.**

| **Gene** | **Symbol^a^** | **Entrez ID** | **Chromosome** | **Start^b^** | **End** |
| --- | --- | --- | --- | --- | --- |
| ENSG00000162551 | *ALPL* | 249 | 1 | 21835858 | 21904905 |
| ENSG00000115211 | *EIF2B4* | 8890 | 2 | 27587219 | 27593353 |
| ENSG00000115234 | *SNX17* | 9784 | 2 | 27593389 | 27599995 |
| ENSG00000163795 | *ZNF513* | 130557 | 2 | 27600098 | 27603657 |
| ENSG00000115241 | *PPM1G* | 5496 | 2 | 27604061 | 27632554 |
| ENSG00000084734 | *GCKR* | 2646 | 2 | 27719709 | 27746554 |
| ENSG00000221843 | *C2orf16* | 84226 | 2 | 27799389 | 27805588 |
| ENSG00000243943 | *ZNF512* | 84450 | 2 | 27805897 | 27858041 |
| ENSG00000176714 | *CCDC121* | 79635 | 2 | 27848506 | 27851879 |
| ENSG00000198522 | *GPN1* | 11321 | 2 | 27851114 | 27874375 |
| ENSG00000119760 | *SUPT7L* | 9913 | 2 | 27873679 | 27886676 |
| ENSG00000163798 | *SLC4A1AP* | 22950 | 2 | 27886338 | 27917840 |
| ENSG00000205334 | *AC074091.13 (LINC01460)* | 100129995 | 2 | 27928653 | 27938599 |
| ENSG00000243147 | *MRPL33* | 9553 | 2 | 27994584 | 28210954 |
| ENSG00000171174 | *RBKS* | 64080 | 2 | 28004231 | 28113965 |
| ENSG00000158019 | *BRE* | 9577 | 2 | 28112808 | 28561768 |
| ENSG00000130561 | *SAG* | 6295 | 2 | 234216462 | 234255701 |
| ENSG00000077044 | *DGKD* | 8527 | 2 | 234263153 | 234380750 |
| ENSG00000085982 | *USP40* | 55230 | 2 | 234384166 | 234475428 |
| ENSG00000118762 | *PKD2* | 5311 | 4 | 88928820 | 88998929 |
| ENSG00000118777 | *ABCG2* | 9429 | 4 | 89011416 | 89152474 |
| ENSG00000165671 | *NSD1* | 64324 | 5 | 176560026 | 176727216 |
| ENSG00000169228 | *RAB24* | 53917 | 5 | 176728199 | 176730745 |
| ENSG00000213347 | *MXD3* | 83463 | 5 | 176728462 | 176739758 |
| ENSG00000169230 | *PRELID1* | 27166 | 5 | 176730775 | 176733960 |
| ENSG00000169223 | *LMAN2* | 10960 | 5 | 176758563 | 176778853 |
| ENSG00000169220 | *RGS14* | 10636 | 5 | 176784838 | 176799602 |
| ENSG00000131183 | *SLC34A1* | 6569 | 5 | 176806236 | 176825849 |
| ENSG00000196570 | *PFN3* | 345456 | 5 | 176827108 | 176827637 |
| ENSG00000131187 | *F12* | 2161 | 5 | 176829141 | 176836577 |
| ENSG00000198055 | *GRK6* | 2870 | 5 | 176830205 | 176869902 |
| ENSG00000164626 | *KCNK5* | 8645 | 6 | 39156749 | 39197226 |
| ENSG00000175003 | *SLC22A1* | 6580 | 6 | 160542821 | 160579750 |
| ENSG00000112499 | *SLC22A2* | 6582 | 6 | 160592093 | 160698670 |
| ENSG00000106049 | *HIBADH* | 11112 | 7 | 27565061 | 27702614 |
| ENSG00000254959 | *INMT-FAM188B* | 100526825 | 7 | 30791753 | 30931696 |
| ENSG00000106125 | *FAM188B* | 84182 | 7 | 30811033 | 30932002 |
| ENSG00000240583 | *AQP1* | 358 | 7 | 30893010 | 30965131 |
| ENSG00000110777 | *POU2AF1* | 5450 | 11 | 111222977 | 111326355 |
| ENSG00000102780 | *DGKH* | 160851 | 13 | 42614176 | 42830714 |
| ENSG00000166415 | *WDR72* | 256764 | 15 | 53805938 | 54055075 |
| ENSG00000169344 | *UMOD* | 7369 | 16 | 20344374 | 20367623 |
| ENSG00000169340 | *PDILT* | 204474 | 16 | 20370492 | 20416059 |
| ENSG00000168447 | *SCNN1B* | 6338 | 16 | 23289552 | 23392620 |
| ENSG00000141376 | *BCAS3* | 54828 | 17 | 58754814 | 59470199 |
| ENSG00000123143 | *PKN1* | 5585 | 19 | 14543865 | 14582679 |
| ENSG00000160951 | *PTGER1* | 5731 | 19 | 14583278 | 14586174 |
| ENSG00000123159 | *GIPC1* | 10755 | 19 | 14588572 | 14606944 |
| ENSG00000019186 | *CYP24A1* | 1591 | 20 | 52769988 | 52790512 |
| ENSG00000100218 | *RTDR1* | 27156 | 22 | 23401593 | 23487208 |
| ENSG00000128266 | *GNAZ* | 2781 | 22 | 23412540 | 23467224 |
| ENSG00000159256 | *MORC3* | 23515 | 21 | 37692487 | 37758446 |
| ENSG00000159259 | *CHAF1B* | 8208 | 21 | 37757676 | 37791313 |
| ENSG00000159261 | *CLDN14* | 23562 | 21 | 37832919 | 37948867 |

^a^54 positionally-mapped genes with unique Entrez IDs and gene symbols

^b^Genomic positions (hg19)

### 2 Figures

#### Figure 1

**A**

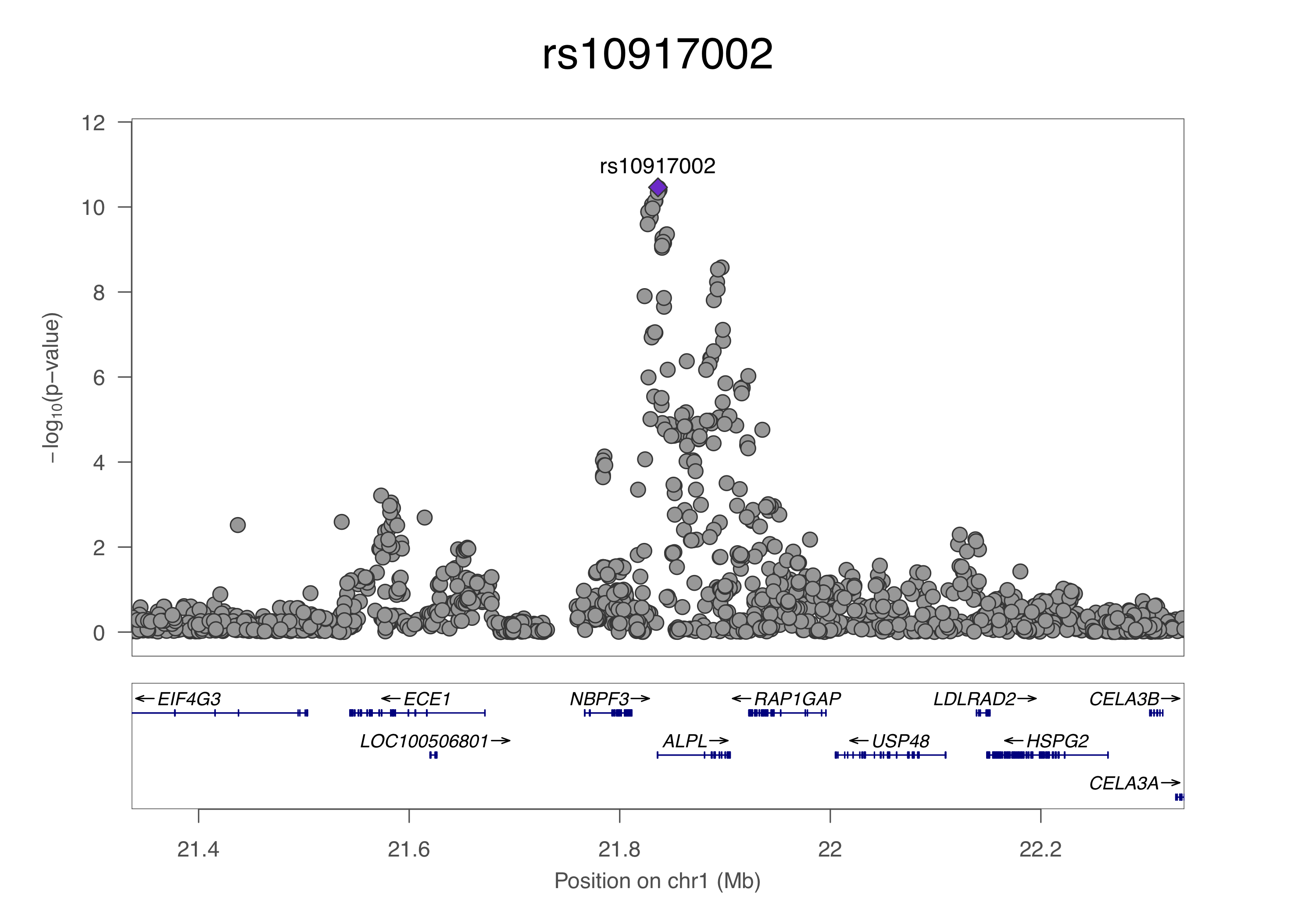

B

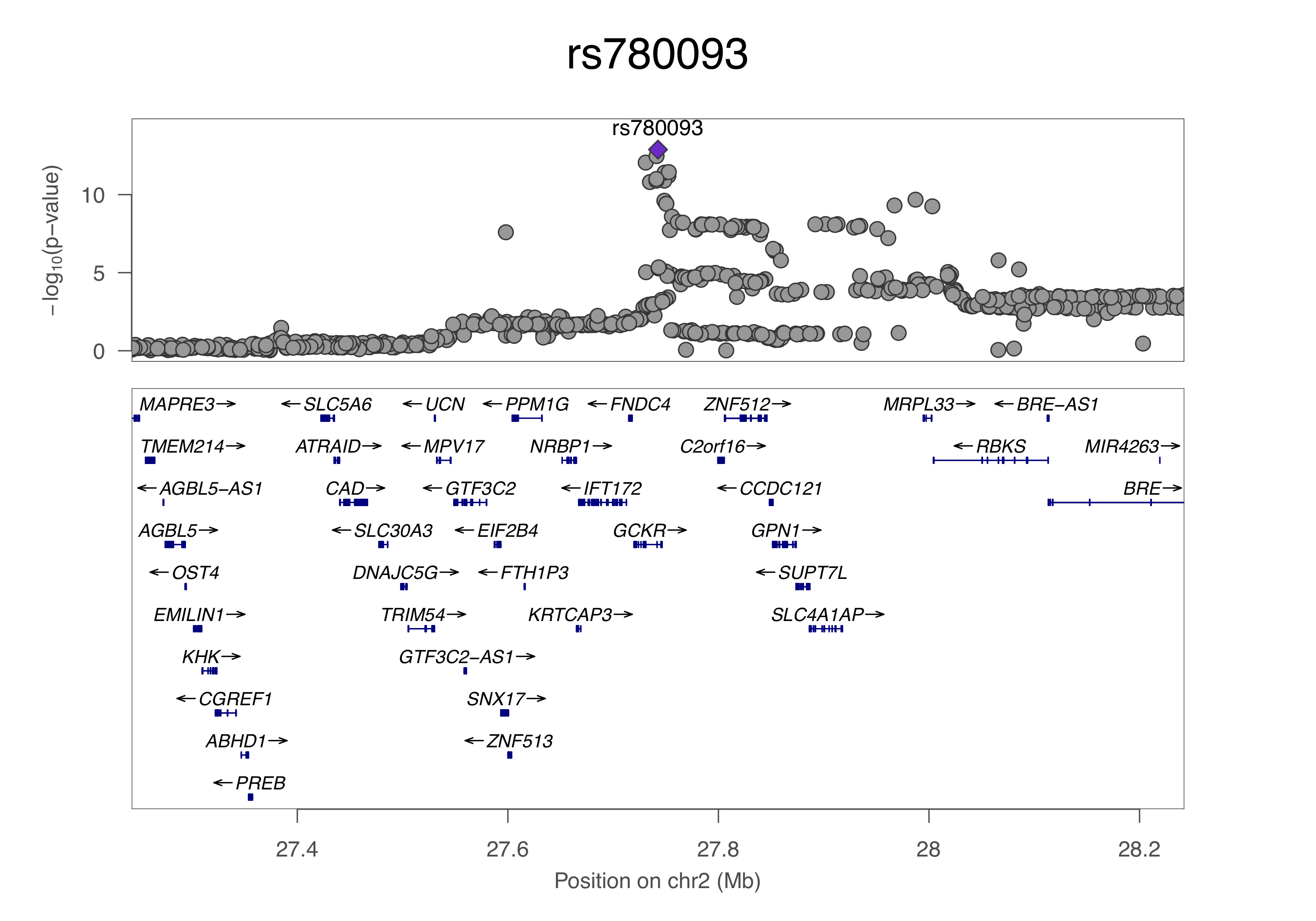

**C**

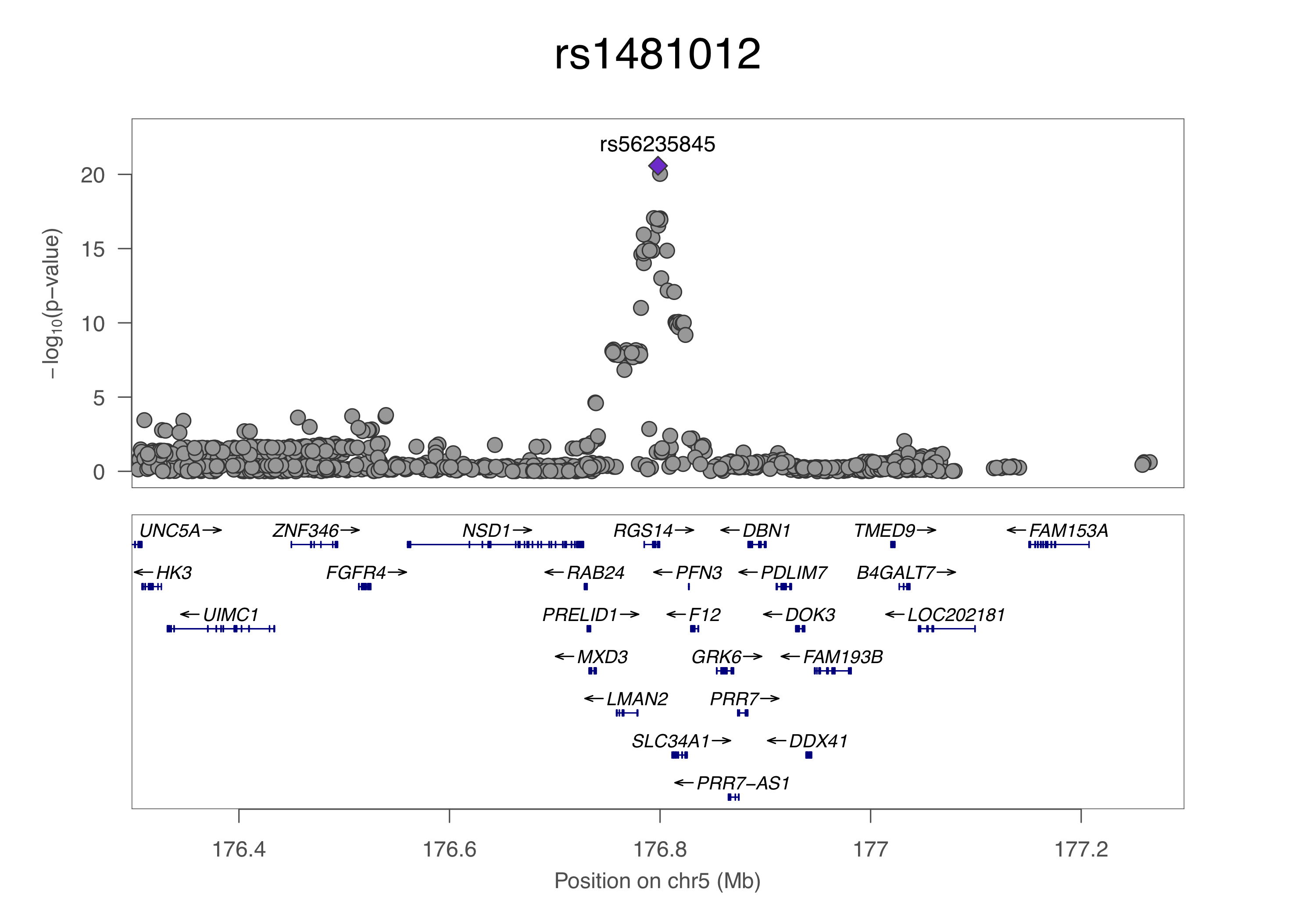

**D**

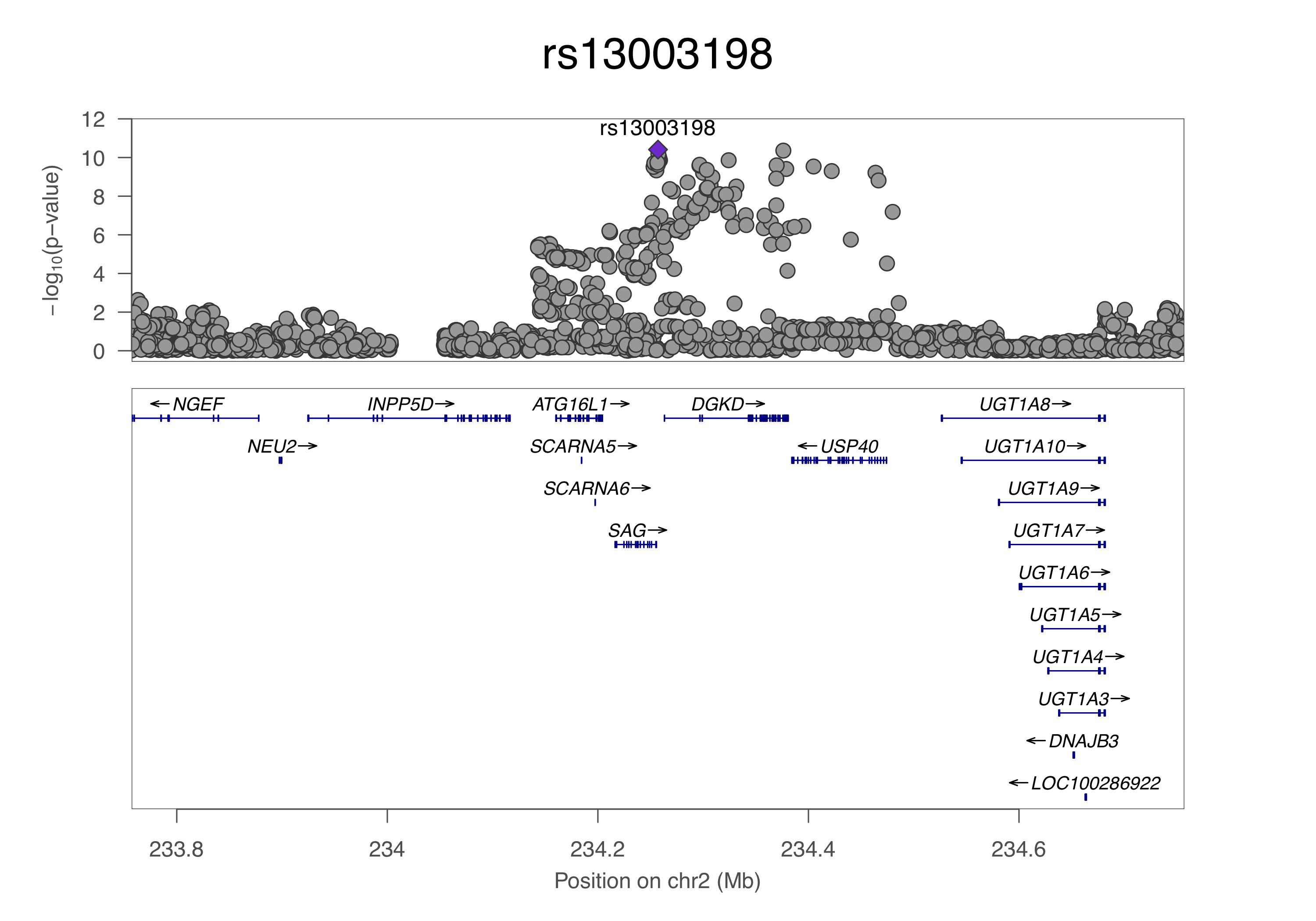

**E**

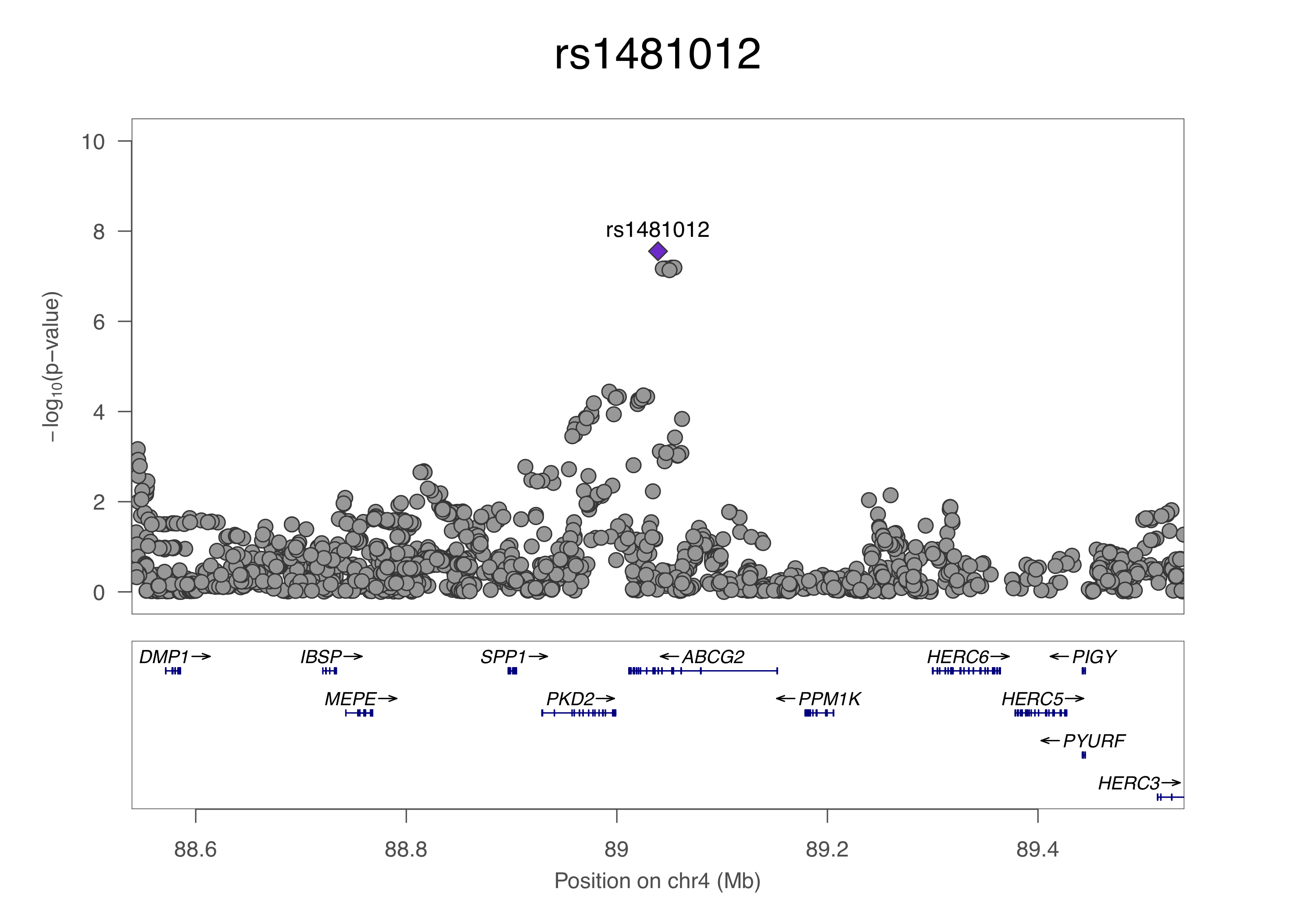

**F**

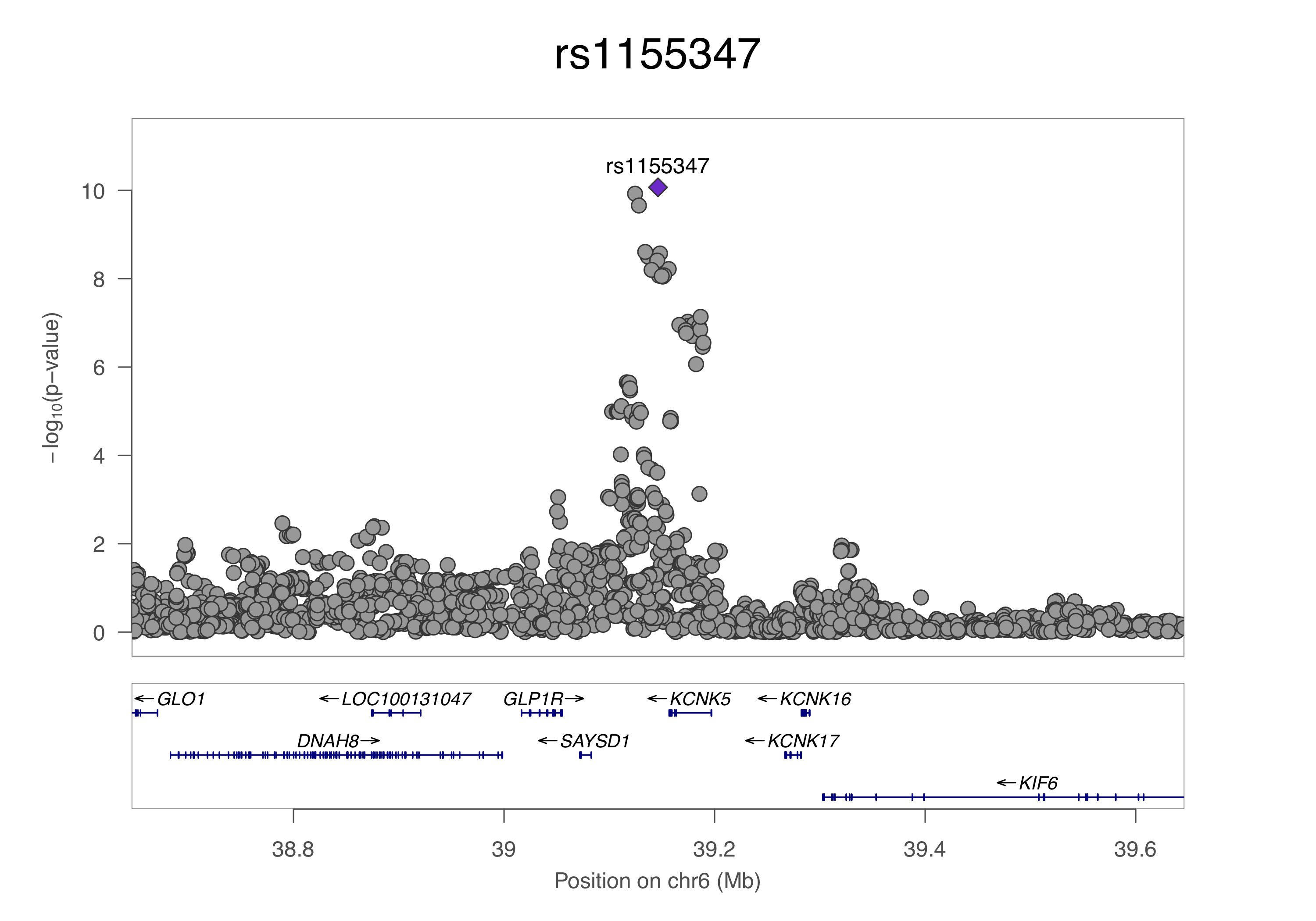

**G**

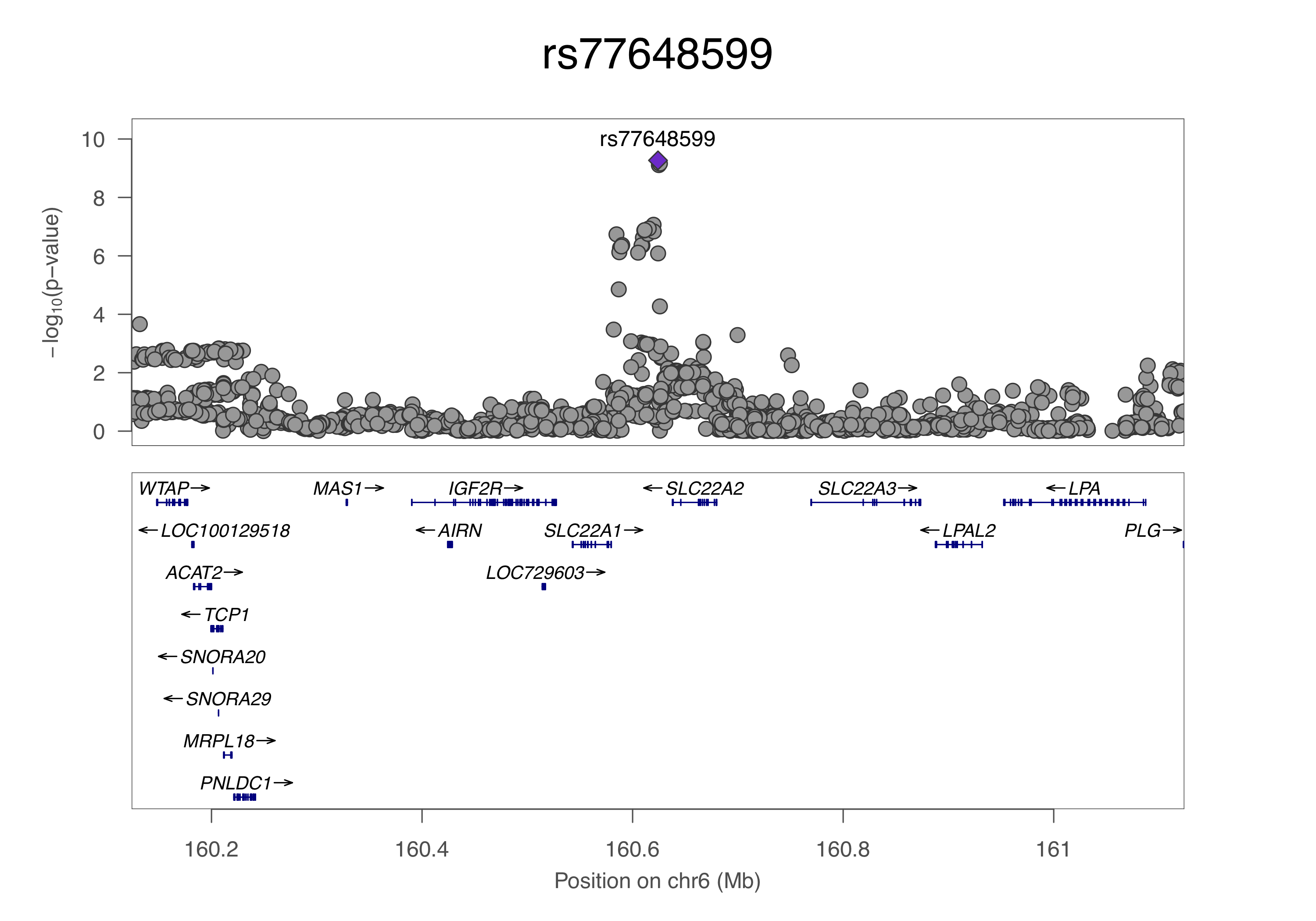

**H**

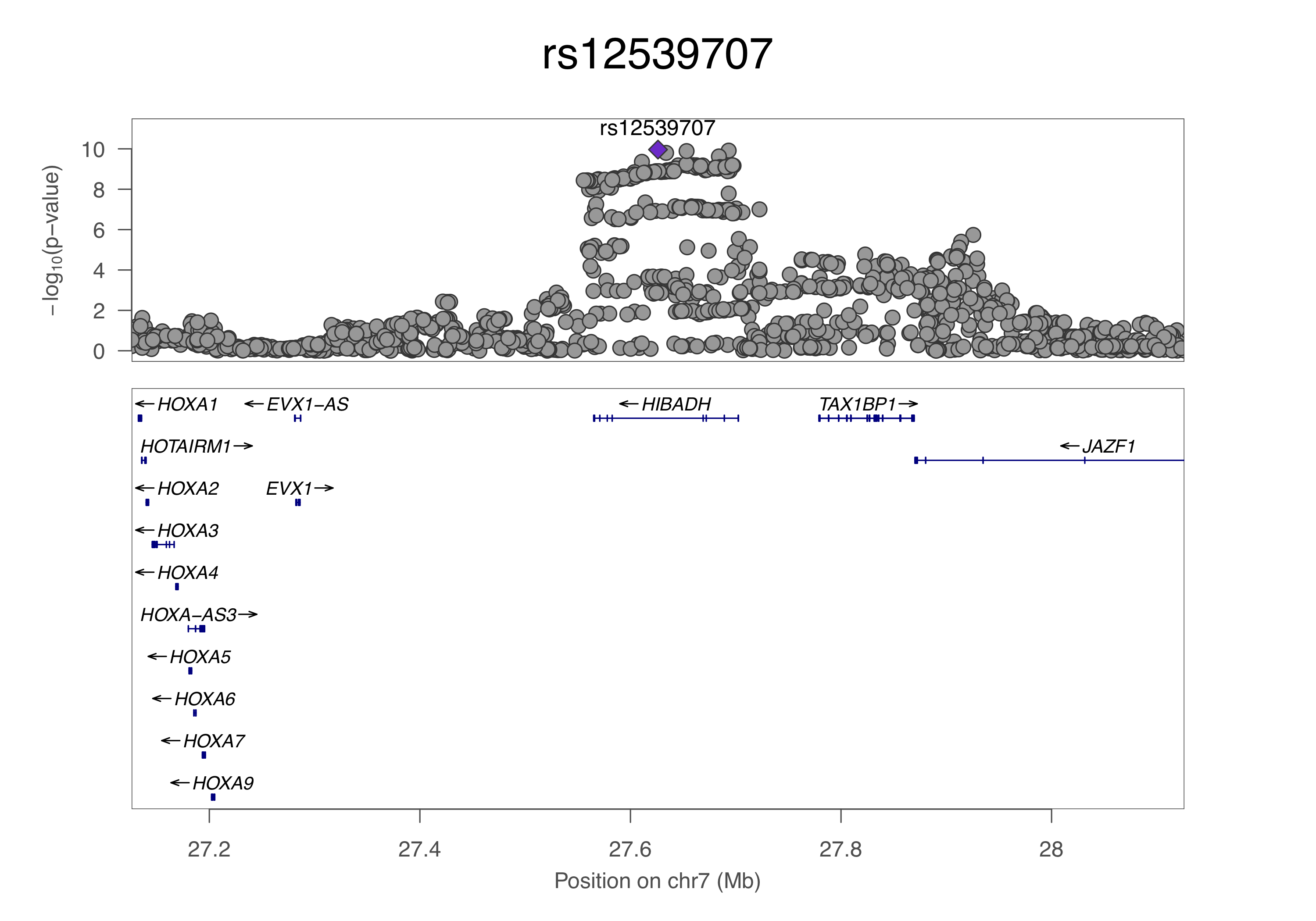

**I**

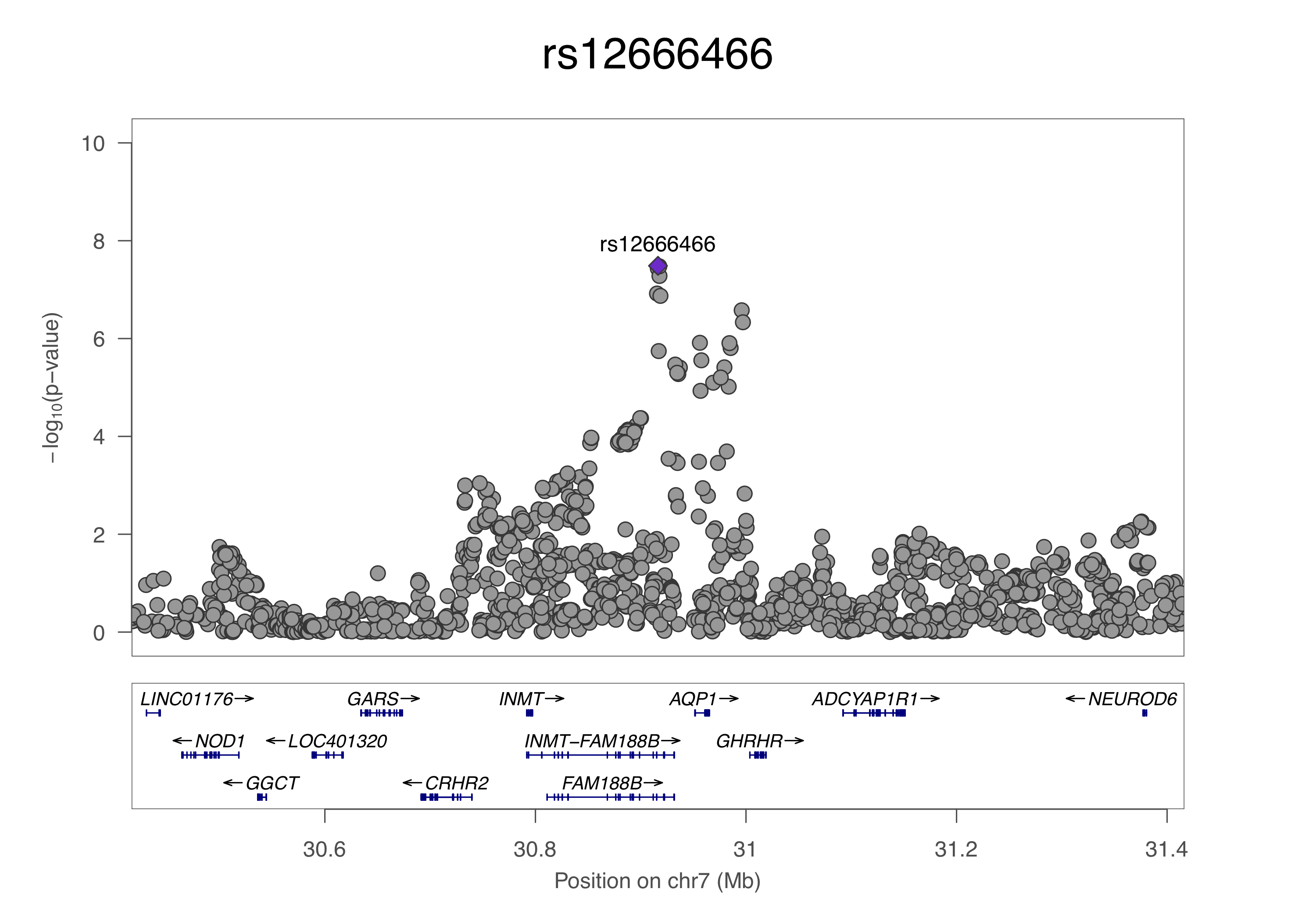

**J**

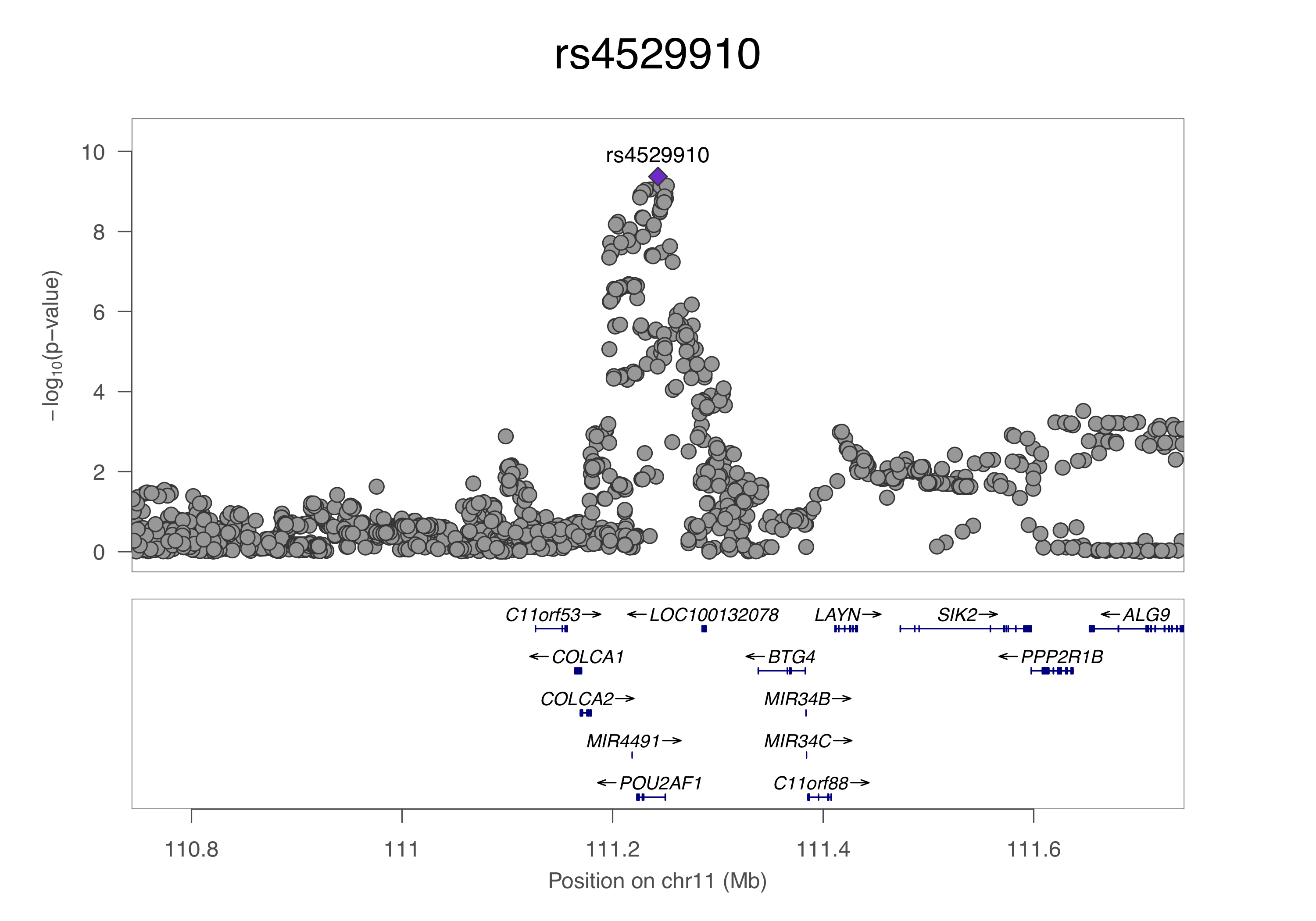

**K**

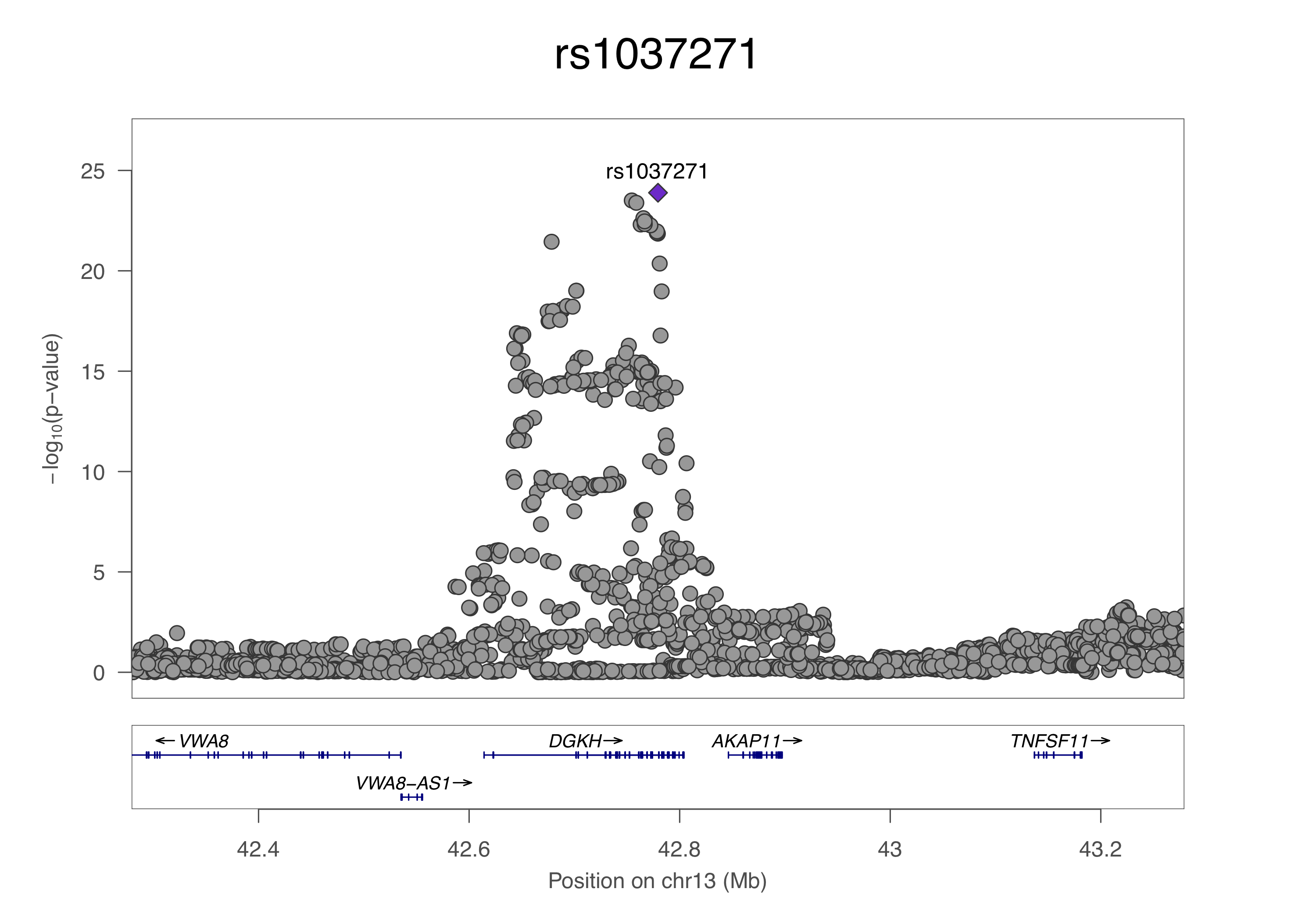

**L**

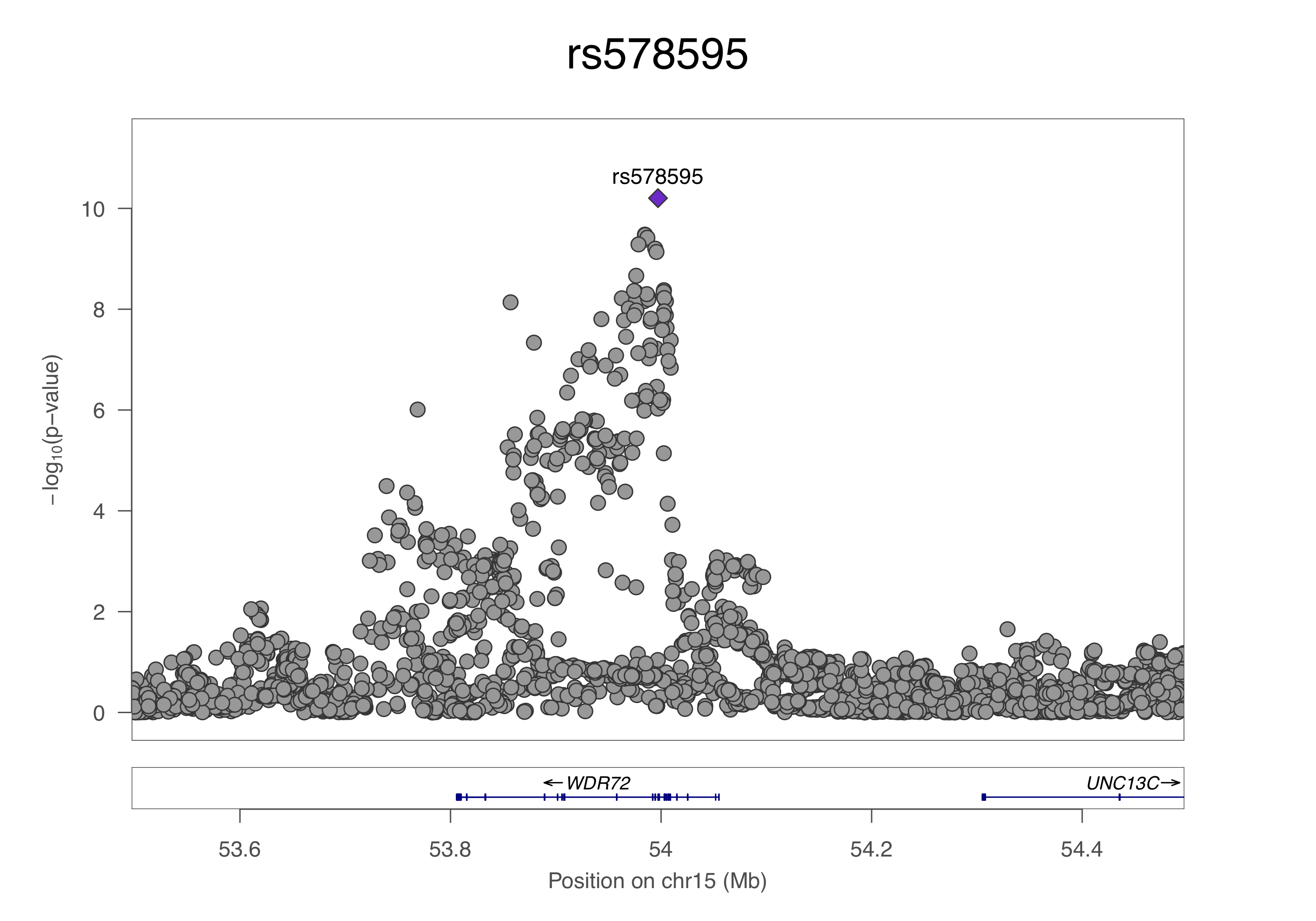

**M**

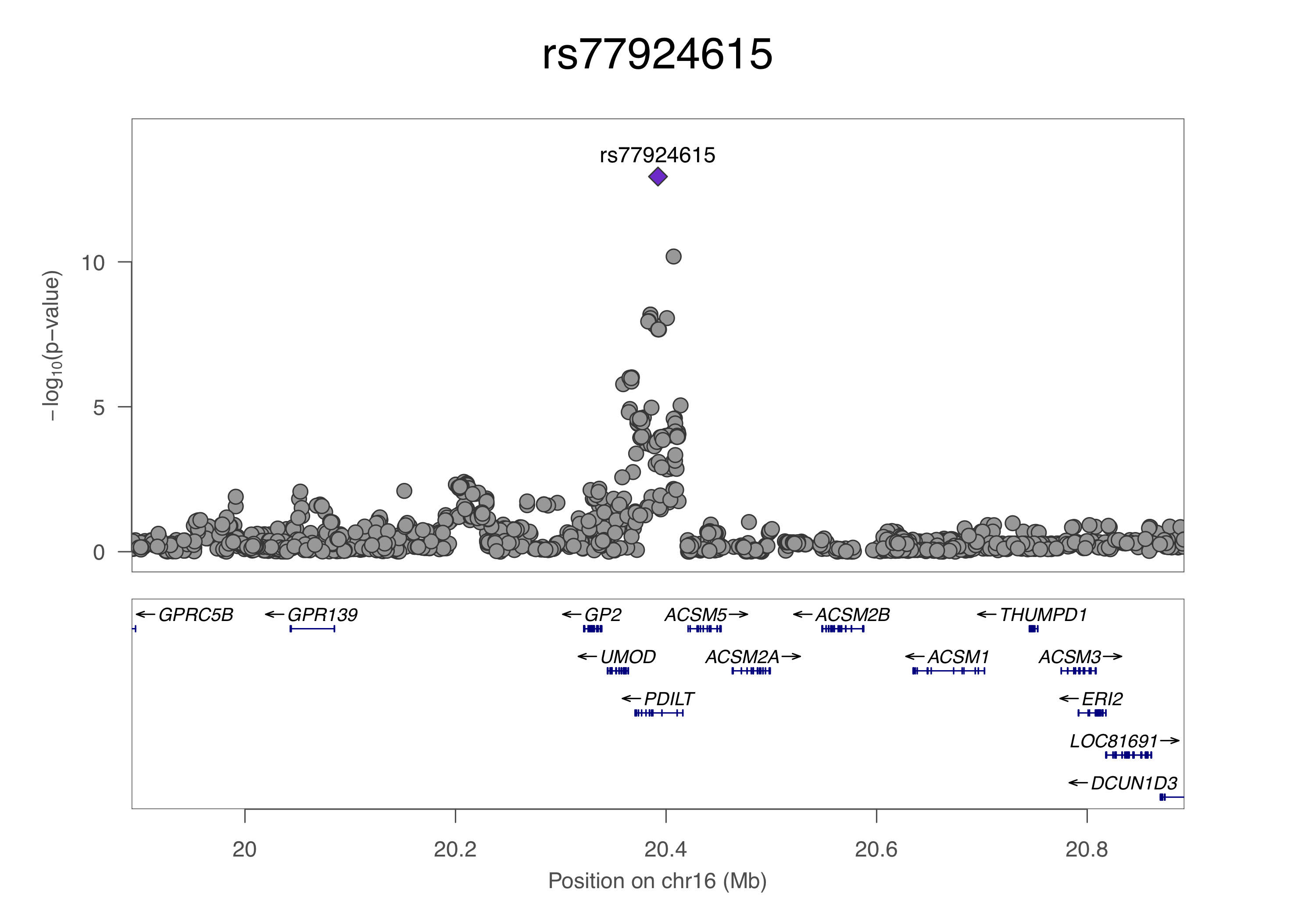

**N**

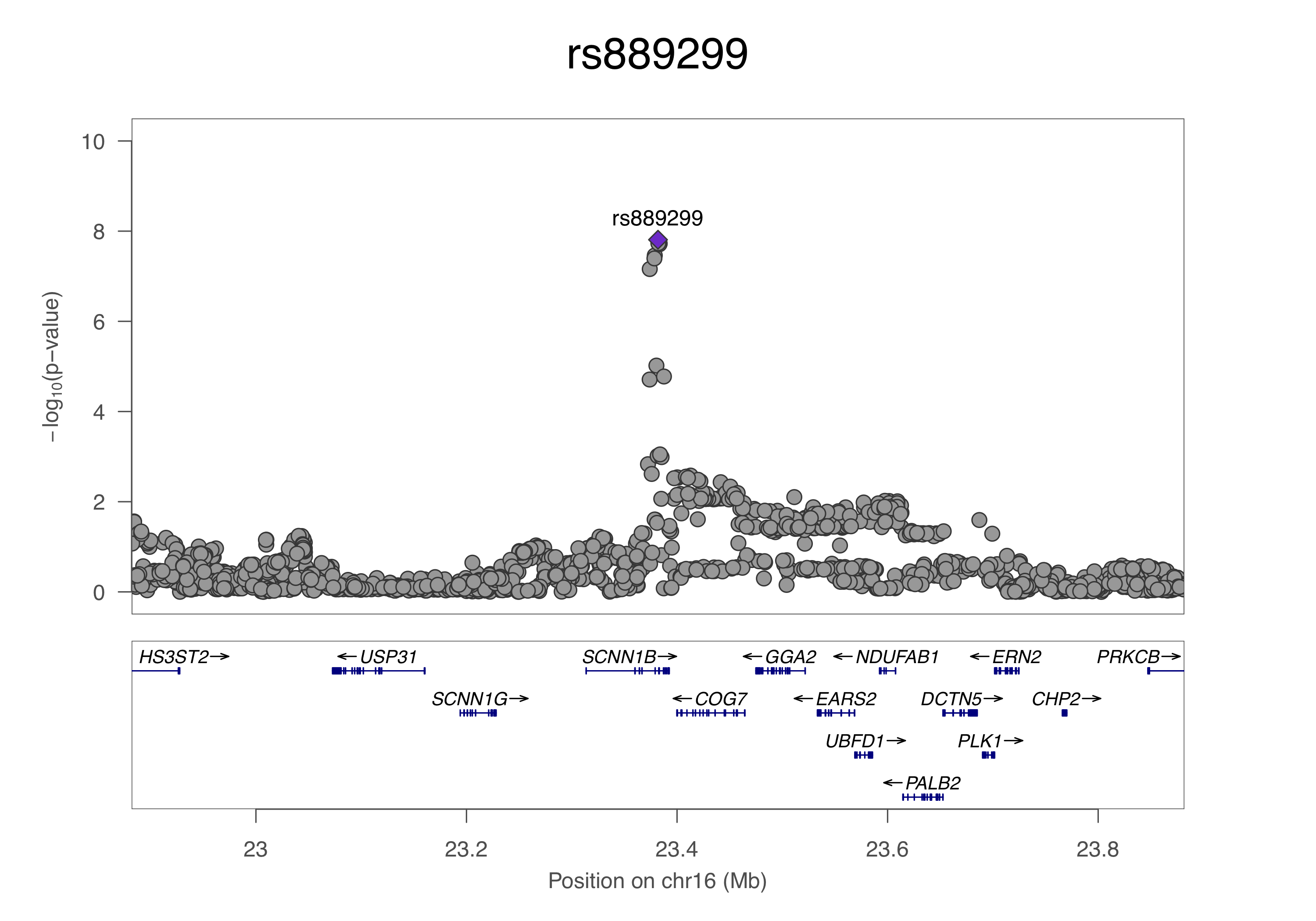

**O**

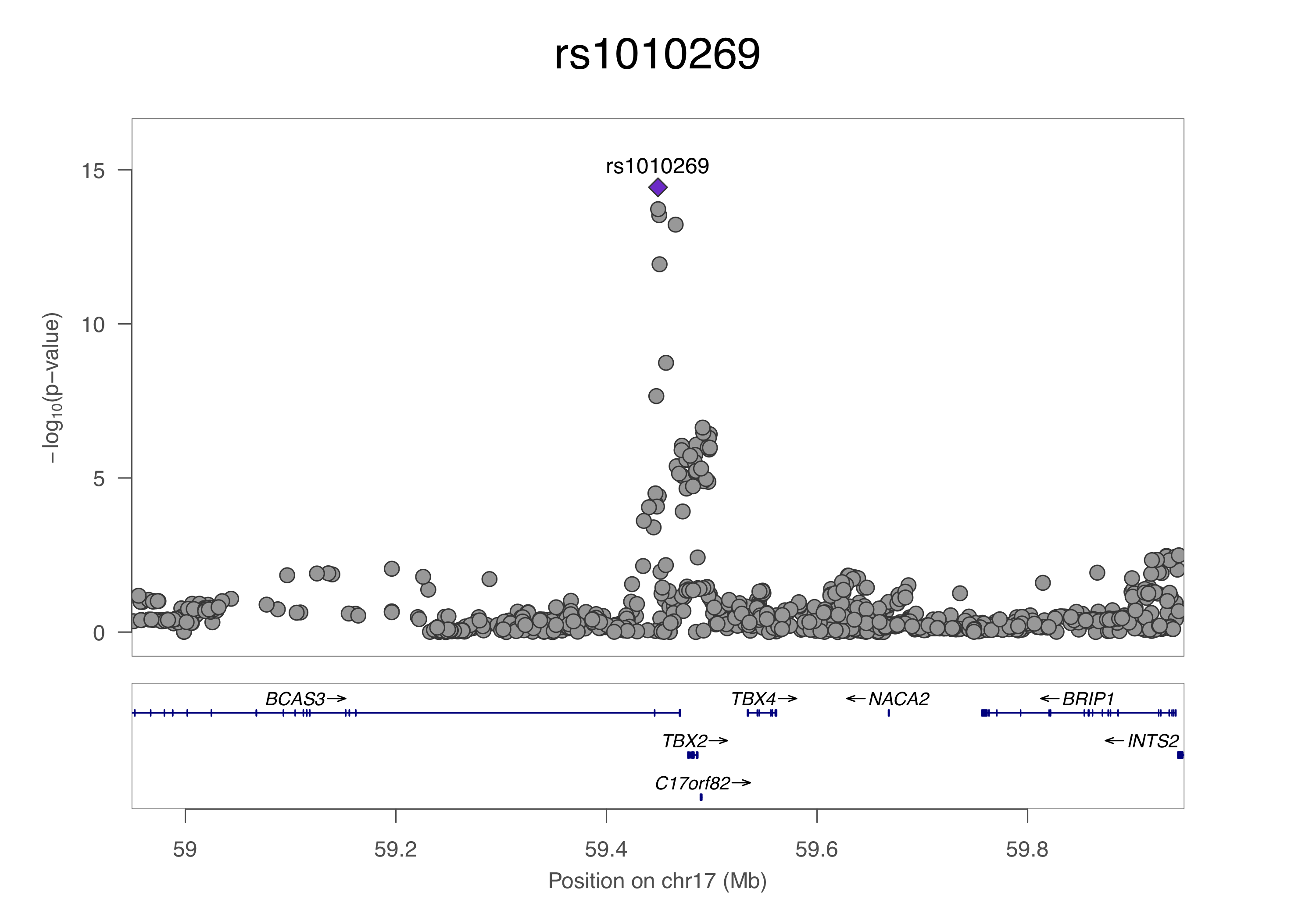

**P**

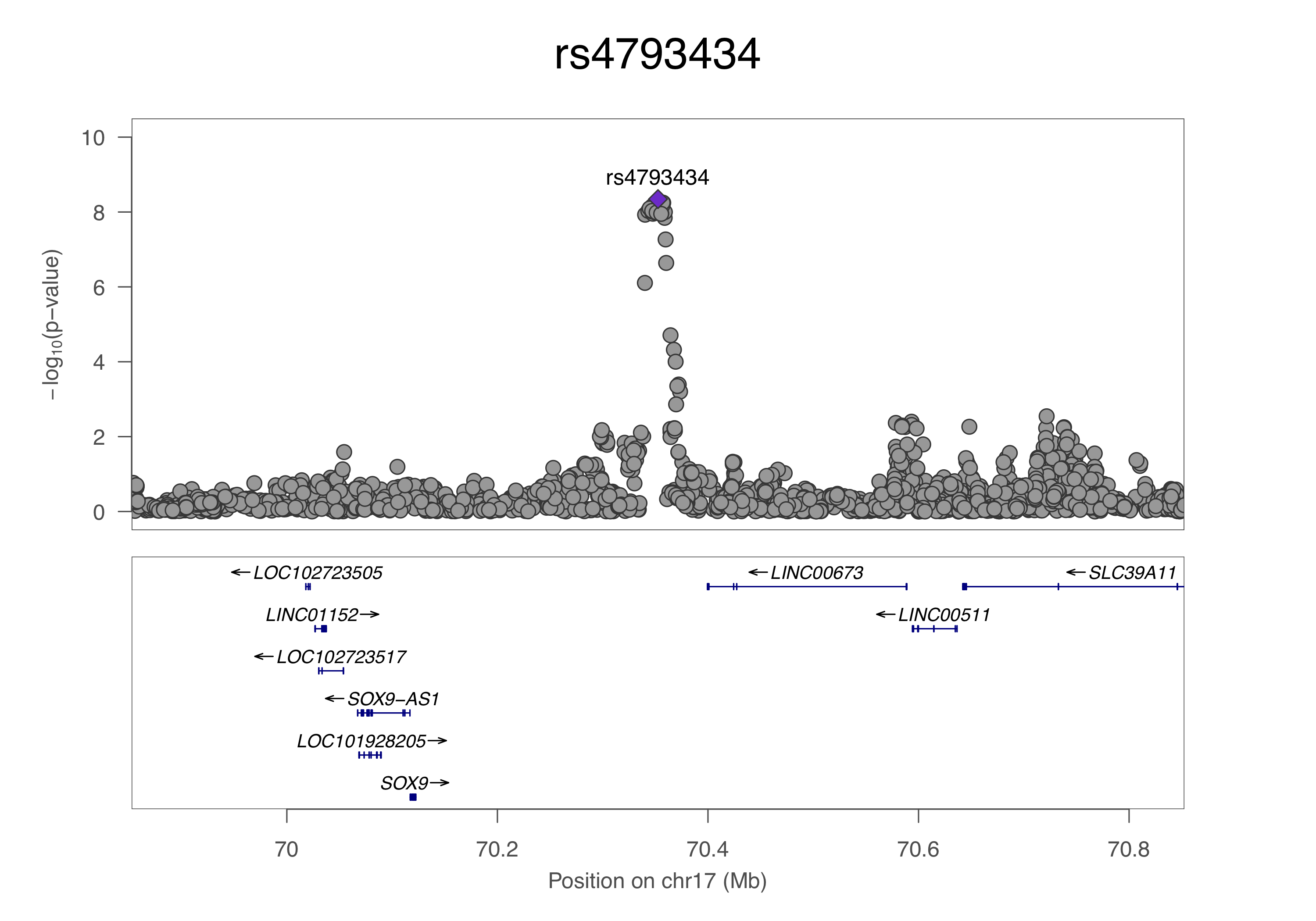

**Q**

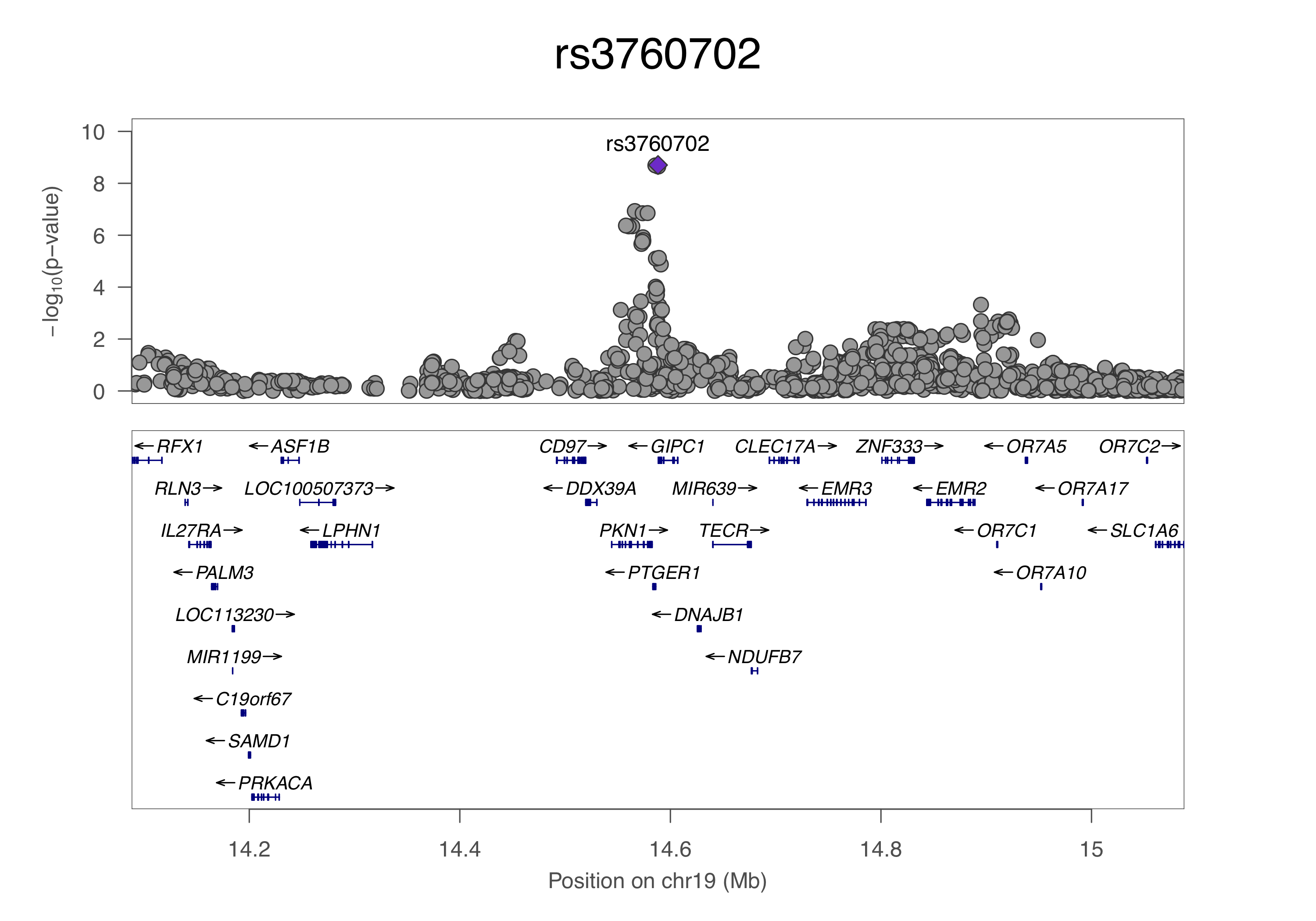

**R**

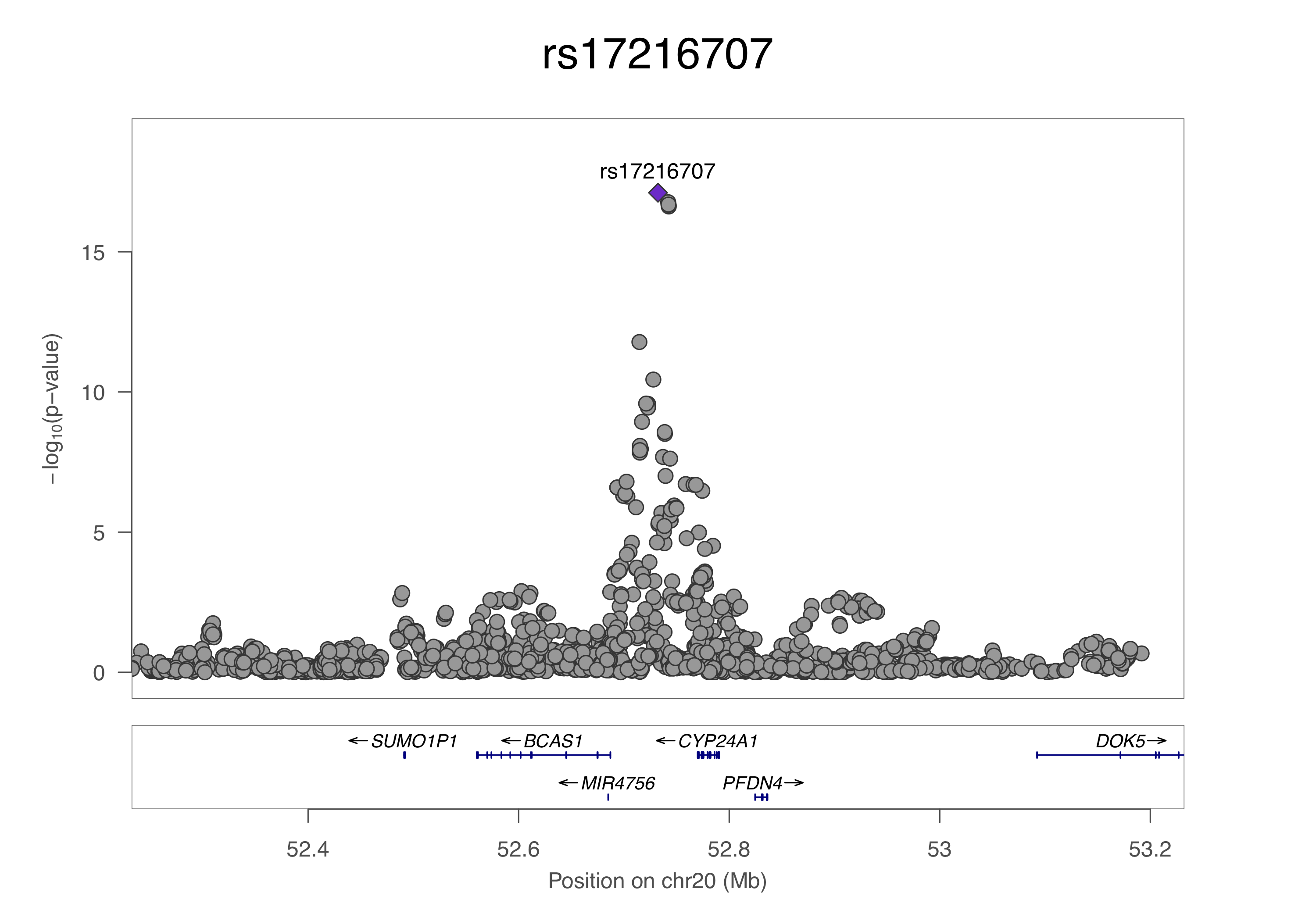

**S**

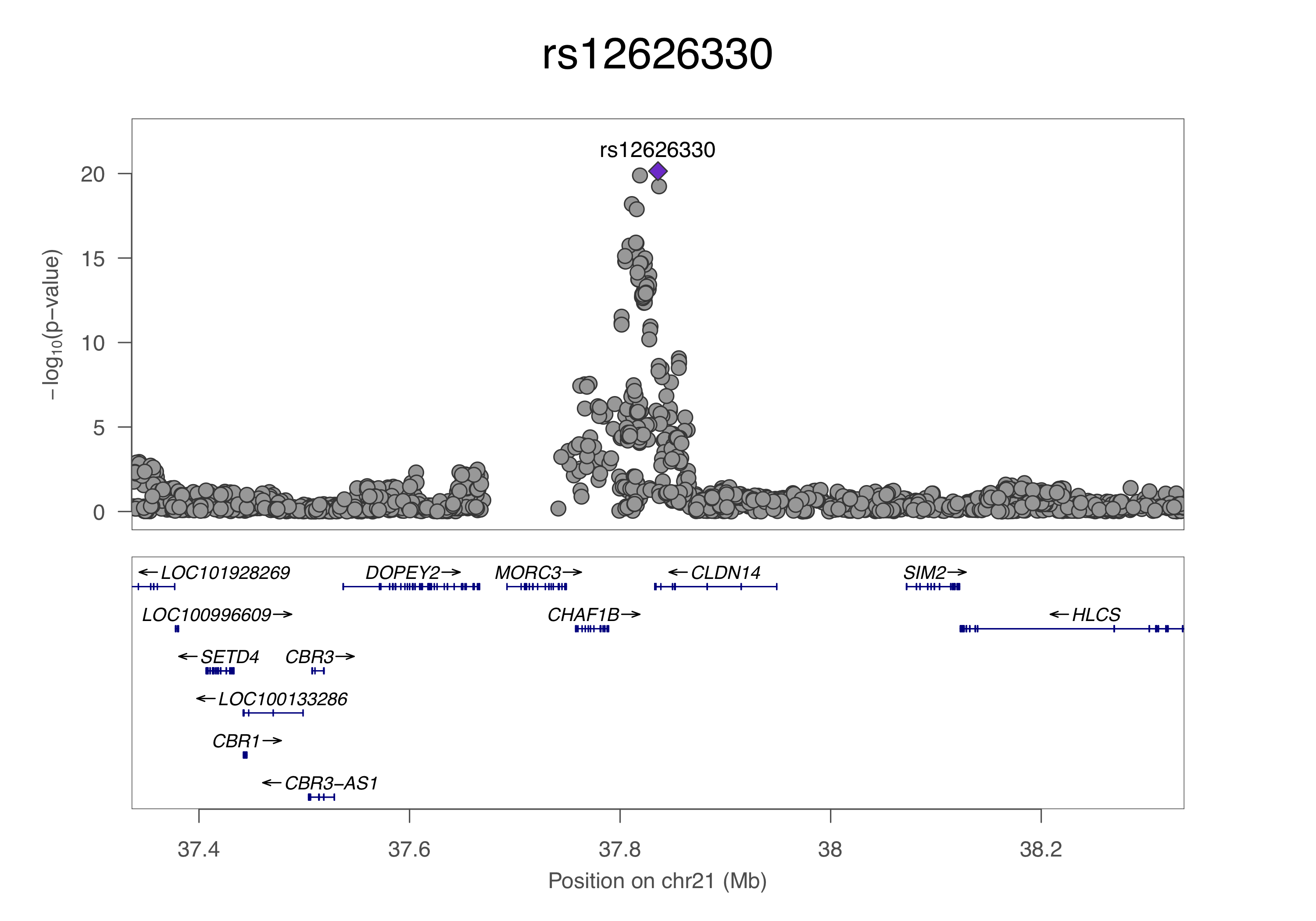

**T**

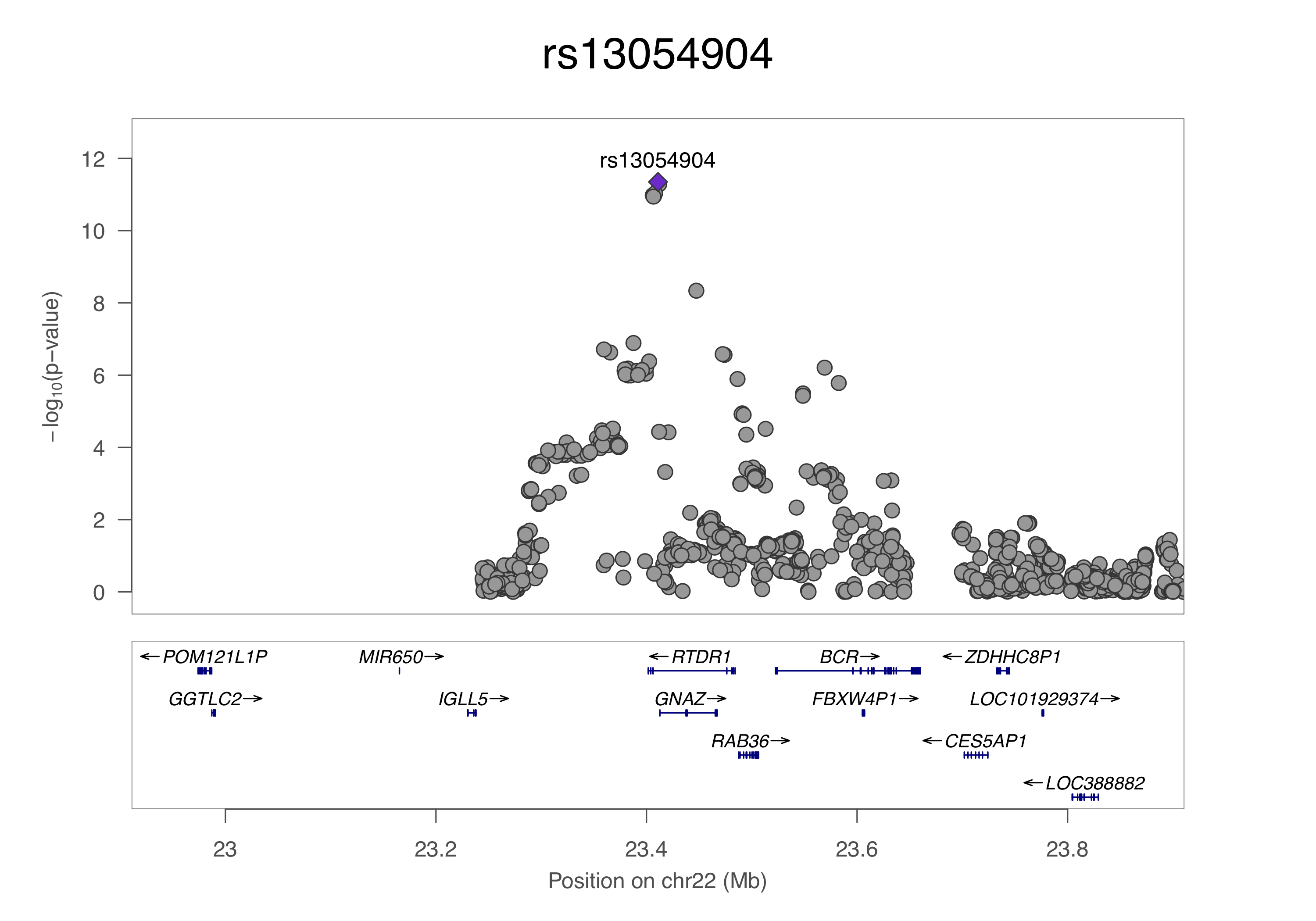

**Supplementary Figure 1: Regional Locus Zoom plots of all GWAS-associated loci in the UK-Japanese meta-analysis.** LocusZoom plots of the 20 top-associated SNPs, ordered by chromosome number and genomic position. SNP position is shown on the x-axis, and strength of association on the y-axis. Genes within 500kb of the index SNP are shown in the lower panel. The position on each chromosome is shown in relation to Human Genome build hg19.

#### Figure 2

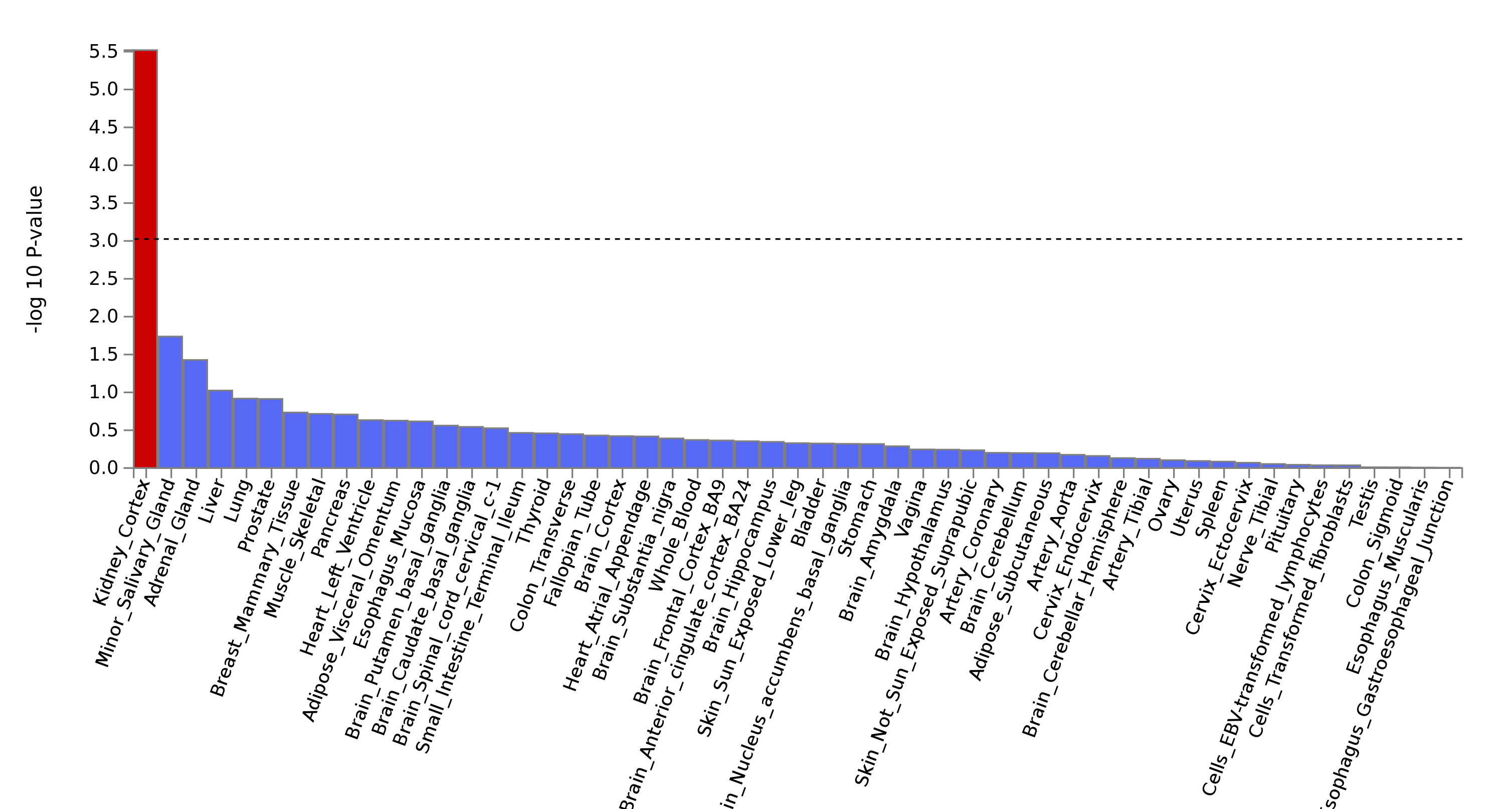

**Supplementary Figure 2:** **Gene-property analysis in MAGMA.** MAGMA Tissue Expression Analysis of GWAS-summary data, implemented in FUMA. This tests the relationship between highly expressed genes in a specific tissue and the genetic associations from the GWAS. Gene-property analysis is performed using average expression of genes per tissue type as a gene covariate. Gene expression values are log2 transformed average RPKM (Read Per Kilobase Per Million) per tissue type after winsorization at 50, and are based on GTEx v6 RNA-Seq data across 53 specific tissue types. The dotted line indicates the Bonferroni-corrected α level, and the tissues that meet this significance threshold are highlighted in red.

#### Figure 3

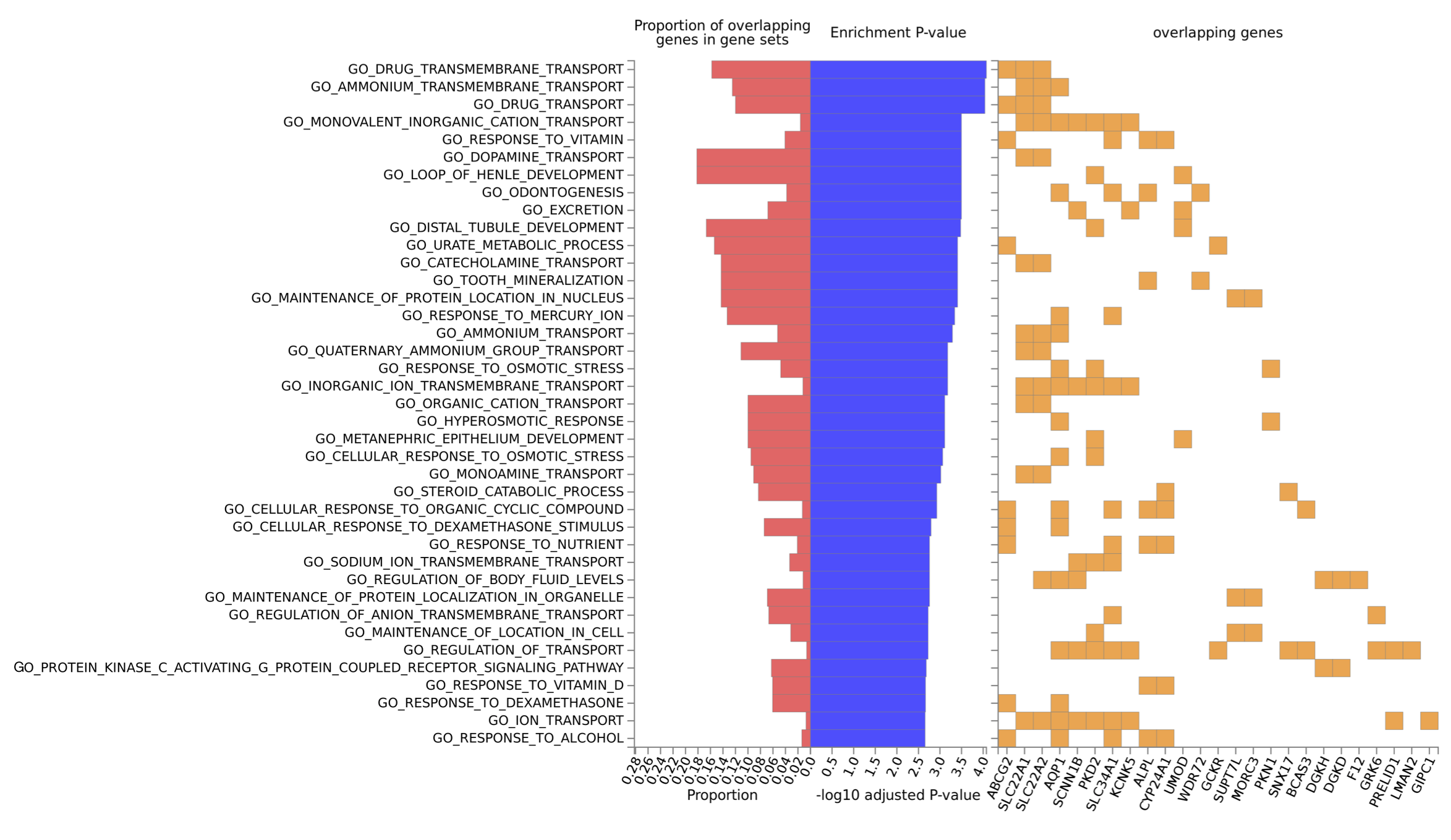

**Supplementary Figure 3: Gene-based enrichment analysis in FUMA.** This analysis was performed using the GENE2FUNC tool in FUMA, using 54 positionally mapped genes with a unique entrez ID and gene symbol. An adjusted enrichment p-value cut-off of p<0.0025 was used. Gene ontologies are ranked by descending enrichment p-value.
